# Short-term Rho-associated kinase inhibitor treatment accelerates primary keratinocyte growth while preserving stem cell characteristics

**DOI:** 10.1101/2022.06.28.497914

**Authors:** Vignesh Jayarajan, George T. Hall, Theodoros Xenakis, Neil Bulstrode, Dale Moulding, Sergi Castellano, Wei-Li Di

**Affiliations:** Infection, Immunity and Inflammation Research & Teaching Department, UCL Great Ormond Street Institute of Child Health, 30 Guilford Street, London WC1N 1EH, UK; Genetics and Genomic Medicine Research & Teaching Department, UCL Great Ormond Street Institute of Child Health, 20 Guilford Street, London WC1N 1DZ, UK; Department of Plastic Surgery, Great Ormond Street Hospital for Children, Great Ormond Street, London WC1N 3JH, UK; Light Microscopy Core Facility, UCL Great Ormond Street Institute of Child Health, 30 Guilford Street, London WC1N 1EH, UK; UCL Genomics, Zayed Centre for Research into Rare Disease in Children, 20 Guilford Street, London WC1N 1DZ, UK

**Author notes:** **Corresponding authors** Professor Wei-Li Di, Infection, Immunity and Inflammation Research & Teaching Department, Molecular and Cellular Immunology, UCL Great Ormond Street Institute of Child Health, 30 Guilford Street, London WC1N 1EH, UK. These authors contributed equally to this work (Vignesh Jayarajan, George T. Hall). These authors contributed equally to this work (Sergi Castellano, Wei-Li Di).

## Abstract

Somatic stem cells can be cultured *in-vitro* and are attractive for cell and gene therapies, but their slow growth in *in-vitro* culture affects survival and stemness and hinders clinical applications. Rho-associated kinase inhibitor (ROCKi) has been used to overcome these obstacles. However, it risks changing the characteristics of stem cells. We found that primary keratinocyte stem cells (KSCs) cultured with the ROCKi Y-27632 for six days exhibited rapid proliferation while maintaining the ability to differentiate. Importantly, after discontinuation of ROCKi treatment, KSC numbers and characteristics were indistinguishable from those in non-treated cultures. We further confirmed that ROCKi treatment resulted in the activation of AKT and ERK pathways, which could support cell survival and proliferation in keratinocytes. We thus concluded that accelerating keratinocyte expansion with short-term ROCKi treatment does not exhaust KSCs’ self-renewal and differentiation capacities, presenting a safe avenue for clinical applications.

## Introduction

Cultured epithelial sheets have been used in autologous epidermal sheet graft therapy for severe burns and chronic ulcers for more than three decades^1^. This approach has recently been combined with gene therapies for rare genetic skin diseases in which gene-corrected epithelial sheets from patients’ keratinocytes, including keratinocyte stem cells (KSCs), are cultured and grafted onto patients with severe skin damage^2-4^. Although other gene therapeutic strategies for genetic skin conditions have been applied, *ex-vivo* gene-modified epithelial sheet gene therapy has been the most successful one^2,5^. The efficient expansion of KSCs in *in-vitro* culture is important for epithelial sheet therapy. However, it remains a challenge to accelerate KSCs growth in primary keratinocyte culture without adverse effects on KSC numbers and self-renewal potential.

Indeed, while human skin biopsies comprise 1–10% of KSCs, only 0.1–1% of KSCs typically survive *in-vitro* cultivation^6,7^. This is because dissociation of KSCs from their surrounding cells and extracellular matrix can trigger apoptotic cell death, known as anoikis^8^. It is important to maintain the KSC population in the culture as only KSCs have self-renewal capacity to give rise to transient-amplifying keratinocytes (TAs) and all other forms of differentiated keratinocytes^9^. There are no unique markers to distinguish these populations, particularly KSCs from TAs, but they can be identified using colony formation assays. Under Green’s keratinocyte culture condition, heterogenous keratinocytes form three types of clones, which are classified as holoclones, meroclones, and paraclones based on their morphology, size, and growth potential^10^. Holoclones have the highest proliferative capacity, with less than 5% aborted colonies upon sub-cultivation, whereas meroclones and paraclones have 5%–95% and >95% aborted colonies upon sub-cultivation, respectively^11^. Holoclone-forming cells, therefore, have hallmarks of stem cells, while meroclone-and paraclone-forming cells contain high proportions of TAs and terminally differentiated cells (TDs). Advances in culture techniques have allowed holoclones to be sub-cultured *in-vitro* to produce a large number of keratinocytes. Epithelial sheets can be formed once these cells attach to each other and stratify^12^. Since epithelial sheet gene therapy requires genetically modified KSCs to produce a large number of gene-corrected keratinocytes for sheet formation, a low yield of holoclones/stem cells due to anoikis in freshly isolated keratinocytes unduly prolongs culture time of epidermal sheets. This leads to the premature exhaustion of KSC proliferative potential, the loss of KSC stemness, resulting in a short-lived epidermal sheet graft gene therapy that hinders its clinical application^5^.

The finding that rho-associated kinase inhibitor (ROCKi) helps evade anoikis-induced apoptosis in cultured human embryonic stem cells is a potential solution^13,14^. ROCKi has been reported to prevent anoikis in embryonic, pluripotent, and prostate stem cells by activating protein kinase B (AKT) and inhibiting caspase 3^13,15-17^. ROCKi can also induce proliferation and growth of hair-follicle stem cells and astrocytes through the activation of extracellular signal regulated kinase (ERK) signalling^18,19^. Therefore, ROCKi has been widely used in somatic stem cell culture, including limbal epithelial cells^20^, pluripotent stem cells^21^, endothelial progenitor cells^22^, and KSCs^23^.

Among the available ROCK inhibitors, the isoquinoline-derivative fasudil and the synthetic compound Y-27632 are the most widely used. Fasudil has been approved in Japan to treat brain haemorrhages^24^ and has also been used in clinical trials on strokes^25^, brain blood flow^26^, angina, and other cardiovascular disorders and ocular diseases^25-27^. ROCK is an effector in the small GTPase Rho pathway. It belongs to the protein kinases A, G, and C family and has pleiotropic functions, including the regulation of cellular contraction, division, polarity, motility, gene expression, and morphology^28^. These functions vary with cell type and status, making it necessary to assess the molecular consequences of ROCKi treatment on various somatic stem cells before it is used clinically. Currently, it is unclear how ROCKi accelerates stem cell growth and whether the ‘forced’ proliferation affects the characteristics of stem cells.

We investigated these questions by culturing human primary keratinocytes including KSCs with Y-27632 (hereafter referred as ‘ROCKi’, unless otherwise indicated). Cultured primary keratinocytes were treated with ROCKi for six days and compared to untreated cells using stem cell markers and single-cell transcriptomics. We then withdrew ROCKi for six days and compared against untreated cells in the same way. In addition, the possible signalling pathways associated with cell proliferation and differentiation following ROCKi treatment were investigated.

## Results

### Six-day ROCKi treatment accelerates colony formation in keratinocytes with effects reversed after withdrawal

Freshly isolated keratinocytes with passage zero (P0) containing heterogenous cell populations including KSCs were cultured in Green’s keratinocyte culture medium with feeder cells and 10µM of ROCKi for six days. The culture media were changed at day three, replenishing with fresh ROCKi. At day six, half of the cells were harvested for evaluation (ROCKi-6D^+^) while the remaining cells were continuously cultivated for further six days without ROCKi (ROCKi-6D^+^6D^−^). Cells without ROCKi treatment but cultured with 0.1% DMSO were harvested at these timepoints and used as negative controls (Control-6D^−^ and Control-6D^−^6D^−^, respectively) (**Figure 1**).

**Figure 1.**
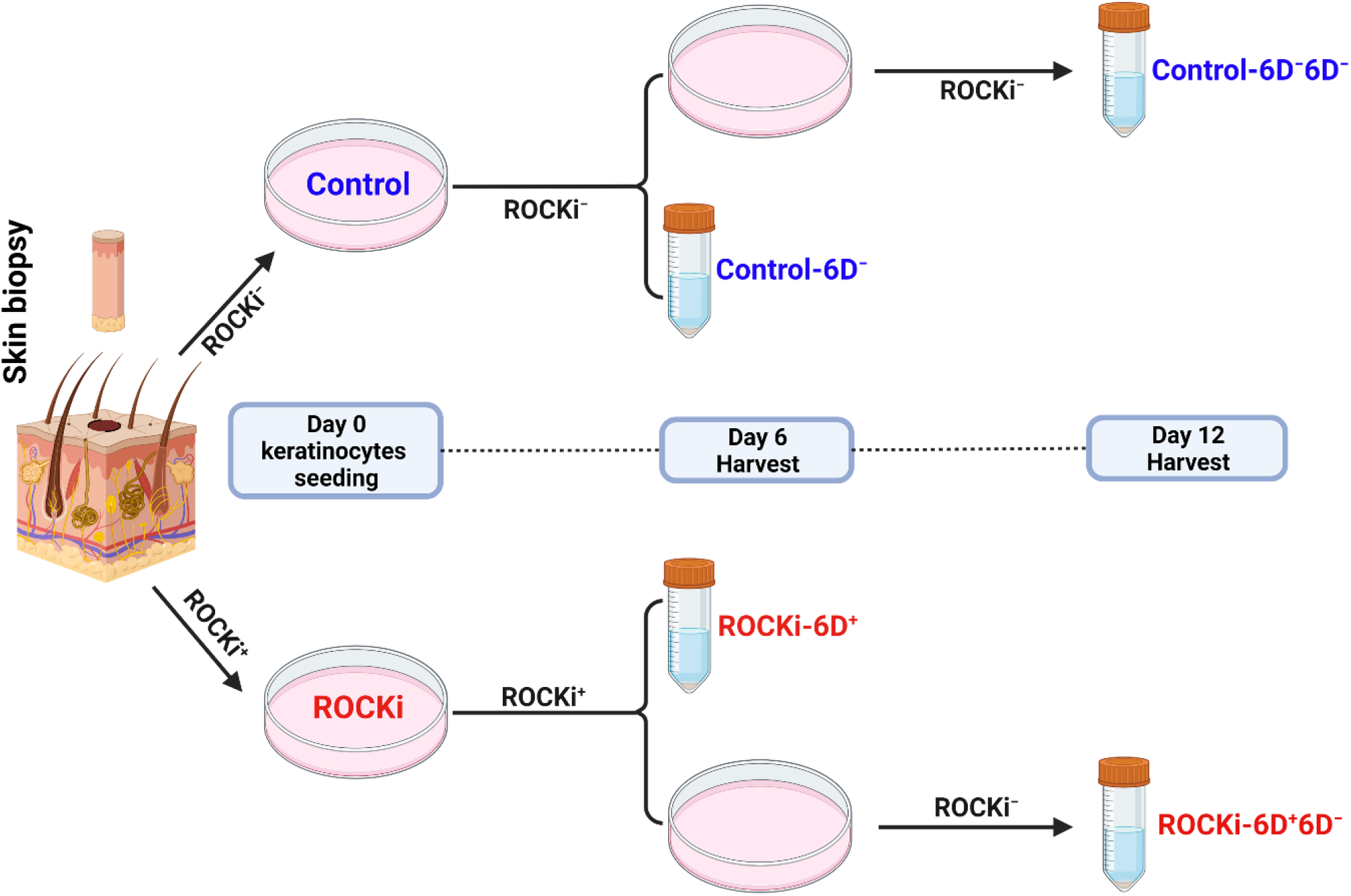
Workflow of experiments and sample collections. Freshly isolated keratinocytes (passage 0) were divided into two populations, either with ROCKi (10µM Y-27632) or without ROCKi treatment (0.1% DMSO) for six days. Half of the cells were harvested on day 6 and the remaining cells were passaged and cultured without ROCKi for additional six days and then harvested.

A 15-fold increase in colony forming efficiency (CFE) and growth rate were detected in ROCKi-6D^+^ cells compared to Control-6D^−^ cells, with the size of ROCKi-6D^+^ cells significantly smaller than those in Control-6D^−^ (**Figure 2A-E**, *p*<0.05). Control-6D^−^ keratinocytes formed tightly packed colonies in the first two days of culture. Conversely, ROCKi-6D^+^ keratinocytes were dispersed with loose connections for the first 2-3 days but at day 4 started to form tightly packed colonies comparable to those in the Control-6D^−^ (**Supplementary Figure S1**). These morphological changes suggested that cells treated with ROCKi were in a proliferating state. Since the mitochondrial content of stem cells is low when they are dormant but increases upon activation and proliferation, mitochondrial mass/content were examined for ROCKi-6D^+^and Control-6D^−^ cells using Mitotracker Green assay. Two distinct cell populations, with low and high mitochondrial mass/content, were observed in both ROCKi-6D^+^ and Control-6D^−^ cells. However, ROCKi-6D^+^ cells had significantly larger populations with high mitochondrial mass than Control-6D^−^ cells (31.84% ± 1.99% v 3.56% ± 3.53%, n = 3, *p*<0 .05) (**Figure 2F** and **G**). Further, cell staining with Mitotracker green showed that cells with high mitochondrial mass were located at the periphery of the colonies (**Supplementary Figure S2A** and **B**). Combining this with the observation that the proliferation marker keratin 14 (K14) was mainly expressed in cells located in the periphery of the colonies (**Supplementary Figure S2C–E**), it strongly suggests elevated activation of stem cells in the culture treated with ROCKi.

**Figure 2.**
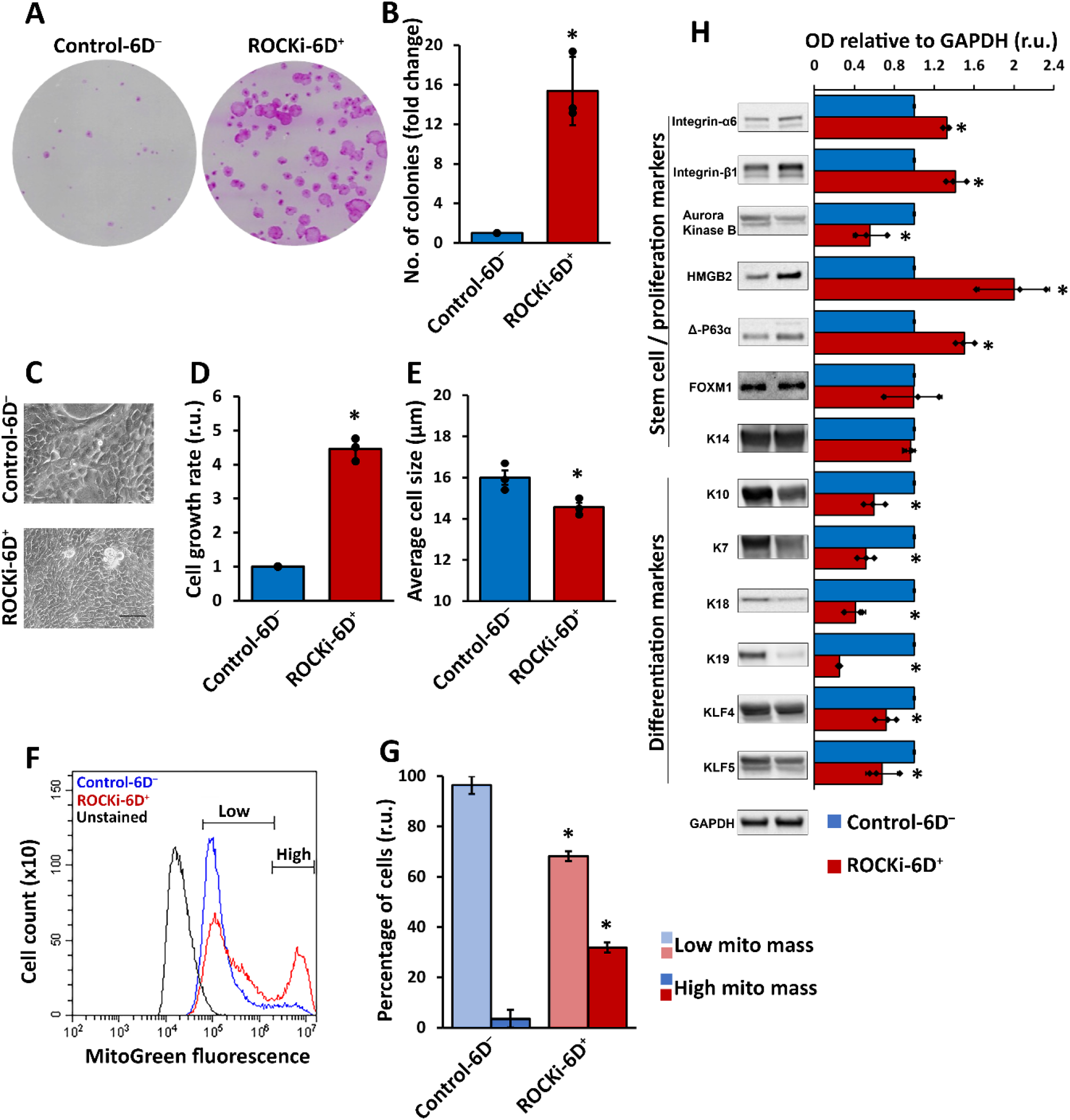
Six-day ROCKi treatment efficiently increased proliferation of primary keratinocytes. Primary keratinocytes at passage 0 were cultured with (ROCKi-6D^+^) and without (Control-6D^−^) ROCKi for six days. Colonies were stained with Rhodanile blue (**A**). A significant increase in colony formation was detected in ROCKi treated cells (ROCKi-6D^+^) compared to non-treated cells (Control-6D^−^) (**B**). The morphology of cultured keratinocytes at day 6. Scale bar = 50μm (**C**). Increased cell growth rate in treated cells compared to non-treated cells (**D**). Decreased average cell size in treated cells compared to non-treated cells (**E**). Histogram of mitochondrial mass in treated and non-treated cells stained with 200nM Mitotracker Green and confirmed by flow cytometry (**F**). Significant increase in cell populations with high mitochondrial mass in treated cells compared to non-treated cells (**G**). Left panel shows the images of representative immunoblots probed with proliferation and differentiation markers (**H**). GAPDH was used as a loading control. The right panel shows the densitometry analysis for immunoblots. Data are presented as mean ± S.D., n=3, * = *p*<0.05.

The expressions of protein markers related to cell proliferation, such as K14, Integrin-α6, Integrin-β1, Aurora Kinase B, HMGB2, FOXM1, Δ-P63α and differentiation including K10, K7, K18, K19, KLF4, and KLF5 in both ROCKi-6D^+^ and Control-6D^−^ cells were examined (**Figure 2H**). Most proliferation protein markers were significantly increased in ROCKi-6D^+^ cells compared to Control-6D^−^ cells, except aurora kinase B which was significantly reduced, and FOXM1 which remained unchanged. In contrast, the expression of differentiation markers was significantly lower in ROCKi-6D^+^ cells compared to Control-6D^−^ cells (**Figure 2H**, n=3, *p*<0.05).

Importantly, in cells which were treated with ROCKi for 6 days and then passaged and continuously cultured for six additional days without ROCKi treatment (ROCKi-6D^+^6D^−^), the colony formation efficiency, cell proliferation rate, cell size and mitochondrial mass in the cells returned to levels comparable to those of non-treated cells (Control-6D^−^6D^−^) (**Figure 3A-G**). The expression of protein markers related to cell proliferation in ROCKi-6D^+^6D^−^ was also reduced to a level similar to that in the Control-6D^−^6D^−^ **(Figure 3H**).

**Figure 3.**
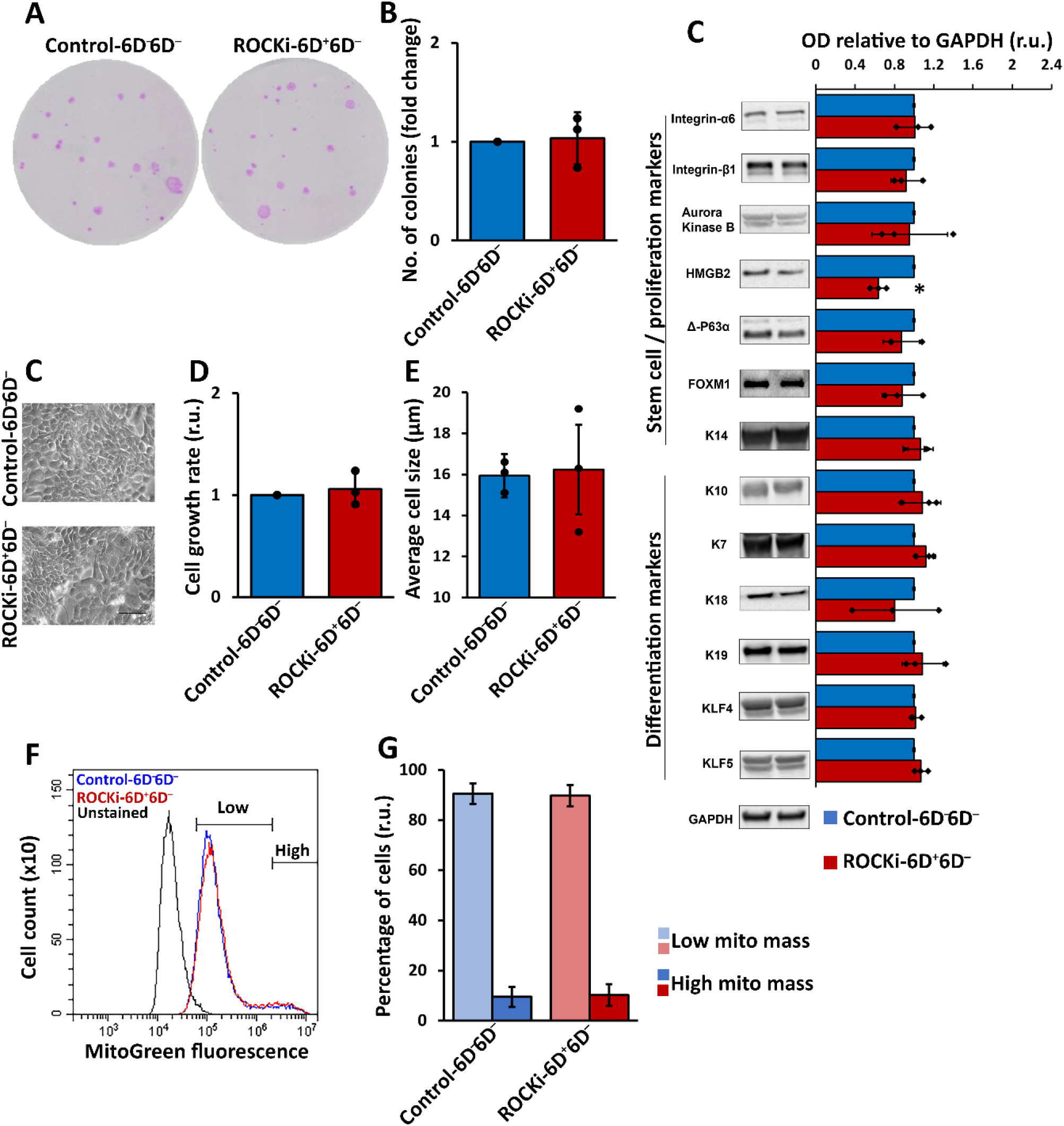
Cells reversed their proliferation status following withdrawal of ROCKi treatment. Control-6D^−^ and ROCKi-6D^+^ cells were passaged once and cultured without ROCKi (Control-6D^−^6D^−^ and ROCKi-6D^+^6D^−^) for additional six days. Colonies were stained with Rhodanile blue (**A**). No significant change in colony formation was detected in treated cells compared to non-treated cells (**B**). The morphology of cultured keratinocytes at day 6. Scale bar = 50μm (**C**). Quantification of the cell growth rate in treated cells compared to non-treated cells (**D**). Quantification of the change in average cell size in treated cells compared to non-treated cells (**E**). Histogram of mitochondrial mass in treated and non-treated cells stained with 200nM Mitotracker Green and confirmed by flow cytometry (**F**). Quantification of the change in cell population with high mitochondrial mass in treated cells compared to non-treated cells (**G**). Left panel shows the images of representative immunoblots with proliferation and differentiation markers (**H**). GAPDH was used as a loading control. The right panel shows the densitometry analysis for immunoblots. Data are presented as mean ± S.D., n=3, * = *p*<0.05.

### Keratinocytes treated with ROCKi increased proliferation and maintained differentiation ability

The proliferation and differentiation potentials in cells with and without ROCKi treatment were evaluated in the organotypic culture *in-vitro*. Fully differentiated multi-epidermal layers – including a basal layer containing cube-shaped basal cells and supra-basal layers containing stratum spinosum, stratum granulosum, and anucleated stratum cornium – were observed in cultures generated using Control-6D^−^, ROCKi-6D^+^, Control-6D^−^6D^−^, and ROCKi-6D^+^6D^−^ cells (**Figure 4A** and **B**). This suggested that ROCKi-induced proliferation did not alter the differentiation ability of the cells. The expression and distribution of protein markers related to proliferation (Δ-P63α) and differentiation (K10, involucrin, and filaggrin) were examined. There were no differences in the expressions and distributions of K10, involucrin and filaggrin (**Figure 4C-J**), but the expression of Δ-P63α increased in ROCKi treated cells (ROCKi-6D^+^) compared to control cells (Control-6D^−^) (**Figure 4C** and **D**). However, it looked no difference of Δ-P63α in the culture following withdrawal of ROCKi treatment (ROCKi-6D^+^6D^−^) compared to non-treated cells (Control-6D^−^6D^−^), indicating that the change of proliferation following a short-term ROCKi treatment was transient (**Figure 4D** and **O**). As Δ-P63α is a marker for keratinocyte stem cells located in the basal layer of epidermis^29^, this staining pattern further suggested that ROCKi might stimulate KSCs from quiescent to proliferative state and thus accelerates KSC differentiation into TAs and, ultimately, TD cells.

**Figure 4.**
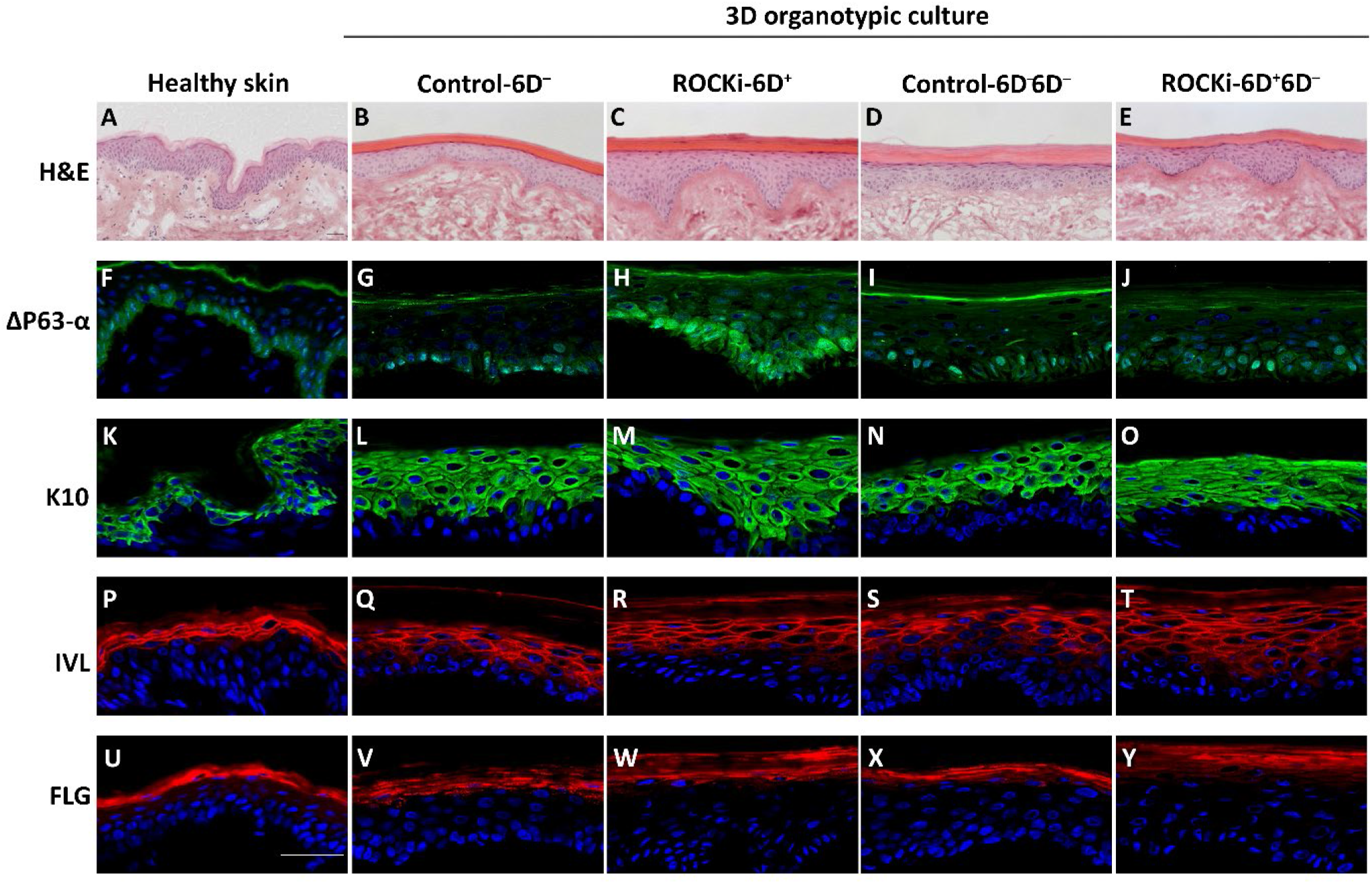
Cells treated with ROCKi for six days had proliferation and differentiation abilities similar to non-treated cells. 3D organotypic cultures were generated using cells with ROCKi treatment (ROCKi-6D^+^),without ROCKi treatment (Control-6D^−^ and Control-6D^−^6D^−^), and after withdrawing ROCKi treatment (ROCKi-6D^+^6D^−^). The 3D culture morphology examined by Haematoxylin and eosin (H&E) staining (**A**–**E)**. Proliferation marker ΔP63-α and differentiation markers K10, involucrin (IVL), and filaggrin (FLG) were examined by immunofluorescence staining (**F**–**Y)**. Increased ΔP63-α expression was detected in ROCKi-6D^+^compared to other groups of cells, but it was returned to the level to non-treated cells after withdrawal of ROCKi treatment. Nuclei were stained by DAPI (blue). Scale bar = 40µM. The expression of these proteins in the skin are shown in **A, F, K, P**, and **U**.

### Reversible changes in the transcriptome of keratinocytes after six-day ROCKi treatment

The transcriptomes of ROCKi-treated and non-treated control cells were compared using single-cell RNA sequencing (scRNAseq). Individual cells that passed the QC criteria were visualised in a uniform manifold approximation and projection (UMAP) plot, where cells group based on the similarity of their transcriptomes. Distinct clusters were observed between cells treated with ROCKi (ROCKi-6D^+^) and non-treated cells (Control-6D^−^) (**Figure 5A**), indicating transcriptomic differences between the groups. In contrast, the transcriptomes became similar in cells following ROCKi’s withdrawal compared to untreated cells (**Figure 5B**), indicating similar transcriptomes. Quantification of overlapping cells clusters using DAseq^30^ corroborated this observation, showing that only 10% of ROCKi-6D^+^ overlapped with Control-6D^−^ (**Figure 5C**), whereas 63% of ROCKi-6D^+^6D^−^ overlapped with Control-6D^−^6D^−^ (**Figure 5D**).

**Figure 5.**
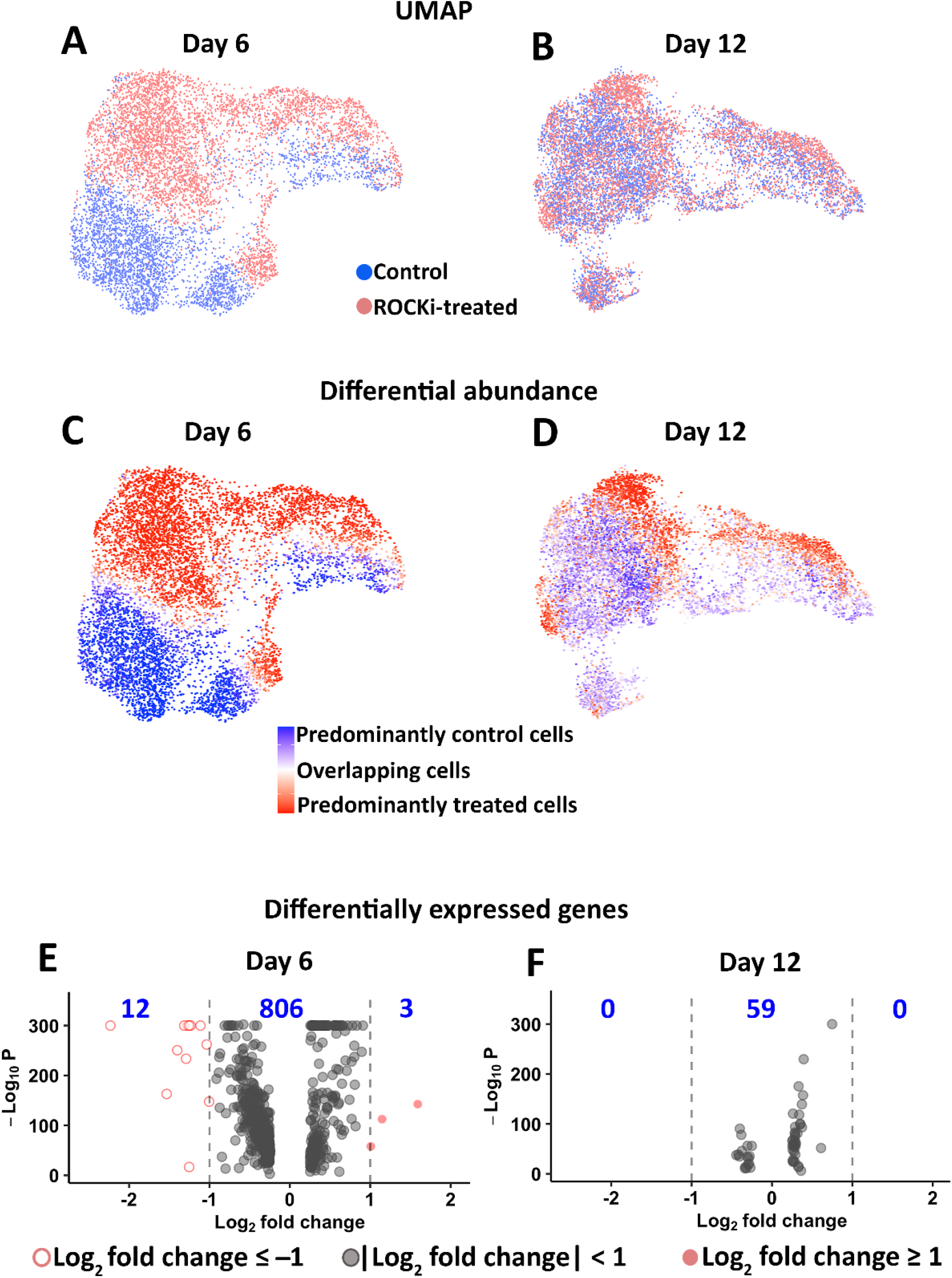
Single-cell transcriptomics: UMAP plots, DAseq analysis, and differential gene expression. UMAP plots were generated and compared between groups at day 6. Distinct clusters between non-treated (Control-6D^−^) and treated (ROCKi-6D^+^) cells were observed (**A**), whereas cells from the ROCKi-withdrawn (ROCKi-6D^+^6D^−^) and non-treated (Control-6D^−^6D^−^) groups clustered together (**B**). Differential abundance analyses verified the overlap observed in (**A**) and (**B**): dark colours indicated little overlap between Control-6D^−^ and ROCKi-6D^+^ cells (**C**) and light colours indicated much overlap between Control-6D^−^6D^−^ and ROCKi-6D^+^6D^−^ (**D**). Volcano plots showing many differentially expressed genes between ROCKi-treated cells (ROCKi-6D^+^) and non-treated cells (Control-6D^−^) (**E**) and far fewer between the ROCKi-withdrawn cells (ROCKi-6D^+^6D^−^) and non-treated cells (Control-6D^−^6D^−^) (**F**). Number of genes in blue indicates significant changes in expression levels.

Fifteen differentially expressed genes with fold changes of at least 2 (i.e. log_2_ fold change >1 or <−1) in ROCKi-6D^+^ compared to Control-6D^−^ cells were identified. Nine genes were downregulated (*KRT7, KRT19, IFI27, DCBLD2, VIM, F3, THBS1, IGFBP7, ARHGAP29, EMP1, FST*) and three genes upregulated (*S100A8, FABP5, KRTDAP*) (**Figure 5E**). *KRT7, KRT19*, and *VIM* have been reported to be involved in cytoskeleton formation and are upregulated in cancer cells^31-33^, while *ARHGAP29* (which codes for Rho GTPase activating protein 29) regulates Rho GTPase signalling by interacting with Rap^34^. The upregulated genes *S100A8, FABP5*, and *KRTDAP* are associated with terminal differentiation of keratinocytes^35,36^. The differentially expressed gene panel suggests that ROCKi might affect cytoskeletal assembly and influence KSCs’ differentiation into TA cells. However, once the treatment was withdrawn, there were no genes with more than 2-fold changes (**Figure 5F**). These results again indicated that withdrawal of ROCKi after a six-day treatment reverted its effects not only at the protein level but also at the gene expression level.

### ROCKi treatment stimulated KSCs to differentiate into transient amplifying keratinocytes

The proportions of KSCs, TAs, and TDs in cultures with and without ROCKi treatment were measured using the expressions of holoclone, meroclone, and paraclone markers described by Enzo, et al. ^37^. ROCKi-6D^+^ had a smaller proportion of KSCs but increased TA and TD cell proportions compared to Control-6D^−^ (**Figure 6A**, 95% confidence interval around total absolute difference in proportions: [0.18, 0.25]). Following withdrawal of ROCKi treatment, the proportions of KSCs, TAs and TDs cells were similar between ROCKi-6D^+^6D^−^ and Control-6D^−^6D^−^ (**Figure 6B**, 95% confidence interval around total absolute difference in proportions: [0, 0.02]). Since these confidence intervals do not overlap, the difference in cell type proportions between treated and control cells was significantly larger during ROCKi treatment than following its withdrawal (*p*<0.05). Further, ROCKi-6D^+^ cells had higher expression of holoclone-specific markers (*AURKB, CCNA2, CKAP2L, FOXM1, HMGB2)*, proliferation markers (*KRT14, TP63, BIRC5*), and early differentiation marker (*KRT10*) compared to Control-6D^−^ cells. However, after withdrawal of ROCKi, the expression of holoclone-specific, proliferation and early differentiation markers was similar in both groups (**Figure 6C**).

**Figure 6.**
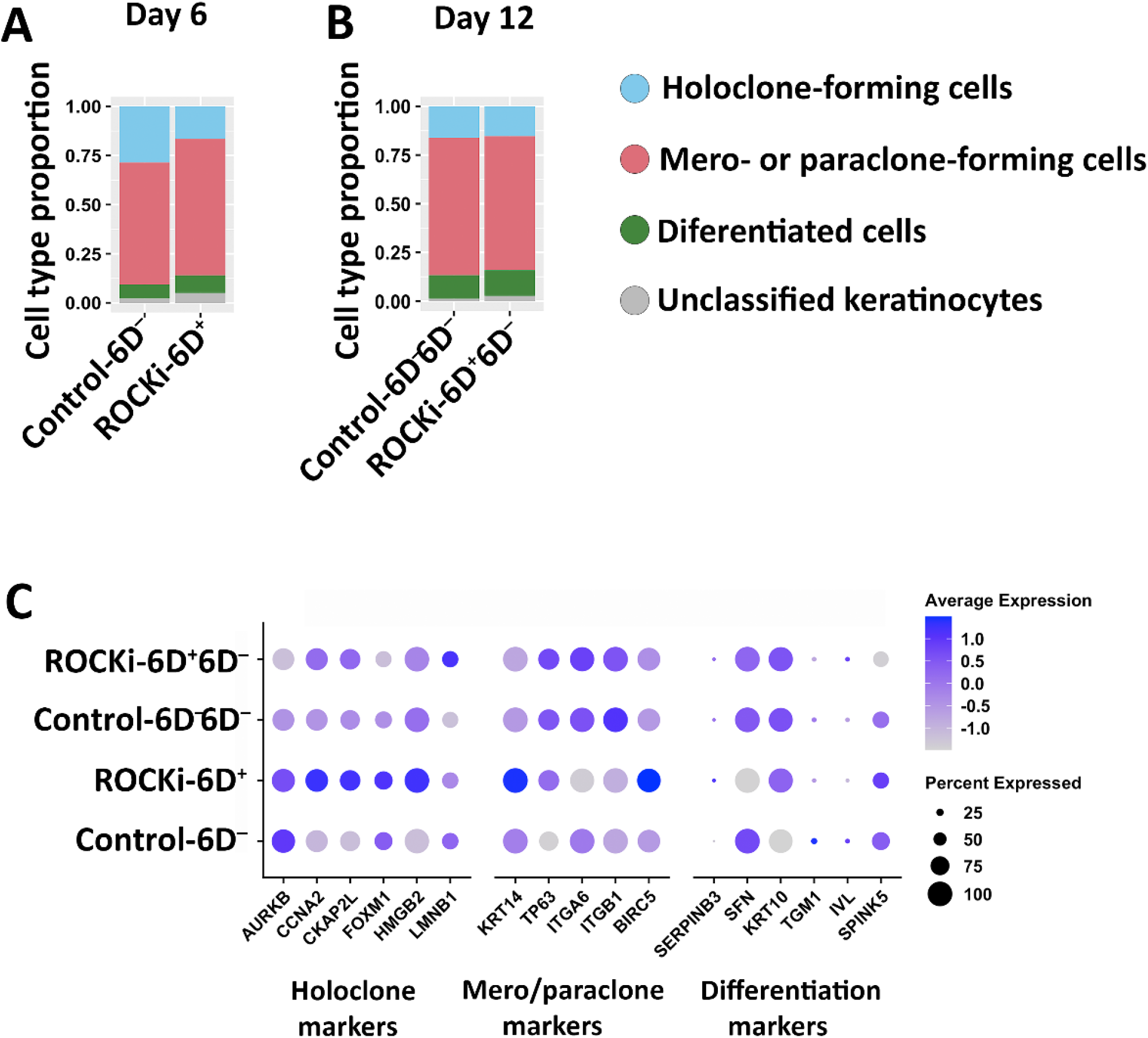
Cell type proportions of cultured primary keratinocytes with and without ROCKi treatment. Cell type populations were analysed using the transcriptome of holoclone, meroclone, and paraclone markers. Differences in cell type proportions was detected between non-treated cells (Control-6D^−^) and ROCKi-treated cells (ROCKi-6D^+^) (**A**), whereas these differences were greatly reduced between non-treated cells (Control-6D^−^6D^−^) and ROCKi-withdrawn cells (ROCKi-6D^+^6D^−^) (**B**). Dotplot showing the expression of markers for holoclone-forming cells, mero-/paraclone-forming cells, and terminally differentiated cells in different samples. The colour of each dot indicates the marker’s expression levels and its size indicates the proportion of cells that express it (**C**).

The differentiation behaviour from KSCs to TAs and finally to TDs was analysed for each sample using single-cell trajectory inference. (**Figure 7A-D**). The inferred pseudotime (differentiation progress) of each cell revealed that ROCKi-6D^+^ cells progressed significantly faster from KSCs to TA then to TD than that in the Control-6D^−^ cells (**Figure 7E**, 95% confidence interval around the difference in cumulative pseudotime distributions: [12.4, 15.5]). However, this acceleration disappeared once ROCKi treatment was discontinued, with the differentiation behaviour of treated cells (ROCKi-6D^+^6D^−^) becoming similar to that seen in untreated ones (Control-6D^−^6D^−^) (**Figure 7E**, 95% confidence interval around the difference in cumulative pseudotime distributions: [1.7, 4.0]). Thus the difference between the differentiation trajectories of treated and control cells is significantly larger during ROCKi treatment than following its withdrawal (*p*<0.05). These results indicate that short-term ROCKi treatment accelerates the speed of differentiation from KSCs into TAs and then to TDs, but this acceleration reverses once ROCKi treatment is discontinued. Importantly, following withdrawal, the numbers of KSCs were also comparable, suggesting that ROCKi did not exhaust KSC self-renewal capacity despite the acceleration of KSC growth during the treatment.

**Figure 7.**
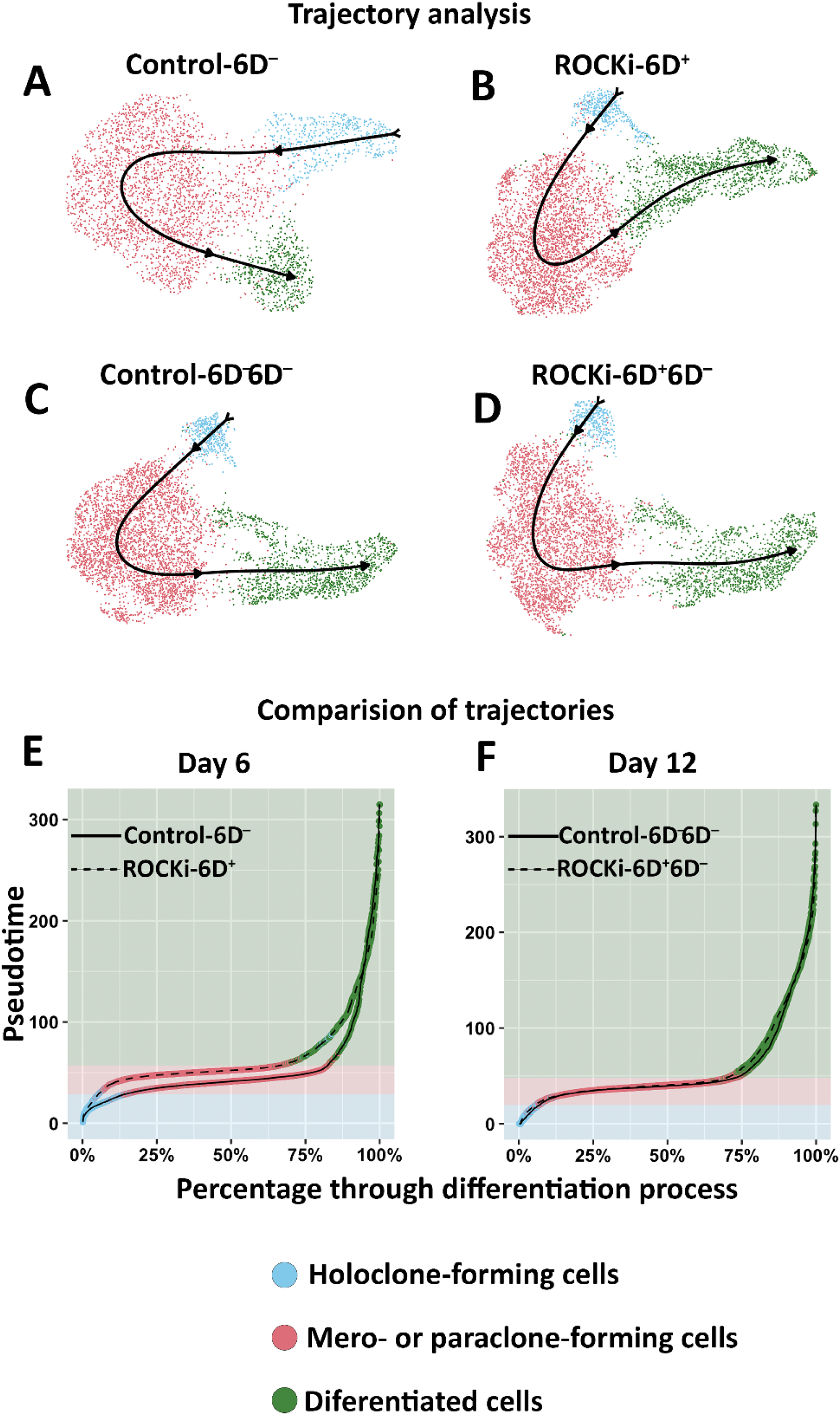
Inferred differentiation trajectories and comparison of pseudotime distributions. Slingshot was used to infer the differentiation trajectories of Control-6D^−^, ROCKi-6D^+^, Control-6D^−^6D^−^, and ROCKi-6D^+^6D^−^ cells, respectively (**A**–**D**). Cell colour corresponds to the three clusters used as input to Slingshot, which correspond to the three cell types. Cumulative distributions of the pseudotimes of Control-6D^−^ and ROCKi-6D^+^ (**E**). Cumulative distributions of the pseudotimes of Control-6D^−^6D^−^ and ROCKi-6D^+^6D^−^ (**F**). The colour surrounding the line in (**E**) and (**F**) corresponds to the cluster of the cell at that point. The shaded portions in (**E**) and (**F**) correspond to the dominant cluster at each stage of differentiation.

### ROCKi treatment rapidly activated AKT1/2 and ERK1/2 signalling

The activation of AKT1/2 and ERK1/2 were investigated to further understand ROCKi’s effects on the cellular signalling related to keratinocyte proliferation. As ROCKi prevents dephosphorylation of myosin light chain (MLC2) by inhibiting the activity of MLC phosphatase^38^, MLC2 phosphorylation in cells treated with ROCKi was assessed. Phosphorylated MLC2 was decreased within 15 minutes and at its lowest 30 minutes post-treatment (**Figure 8A** and **B**). Both AKT1 and AKT2 were activated with a significant increase in phosphorylated AKT1 (Ser473) and AKT2 (Ser474) within 15 minutes of treatment (**Figure 8A** and **C**). Further, the ERK1/2 signalling pathway – including the phosphorylated states of c-RAF (Ser338), MEK1/2 (Ser217/221), and ERK1/2 (Thr202/204) – were assessed. Cells treated with ROCKi significantly increased the phosphorylation of c-RAF and MEK1/2 in 15 minutes. However, the level of phosphorylation of ERK1/2 did not change **(Figure 8A** and **D**). These results indicate that ROCKi acts on primary human keratinocytes by activating AKT1/2 and ERK1/2 signalling pathways.

**Figure 8.**
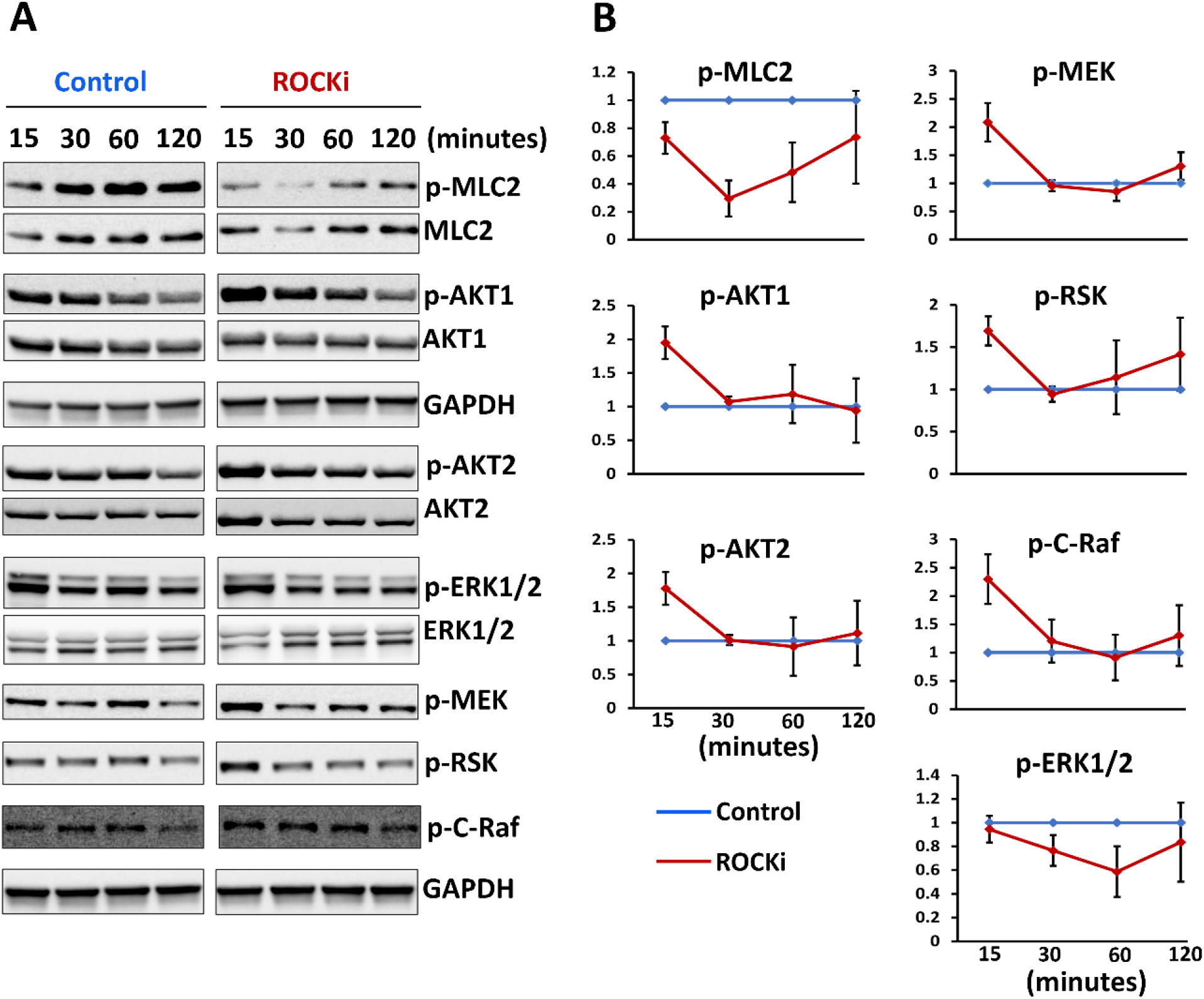
ROCKi activated AKT and MAPK signalling. Representative immunoblot images show the changes of phosphorylation of kinases involved in AKT and MAPK signalling in cells treated with ROCKi at different time points (**A**). GAPDH was used as a loading control. Densitometry quantification of immunoblot at different time points (**B**). Data were normalized to GAPDH and are presented as mean ± S.D., n=3.

## Discussion

Epithelial sheet gene therapies require gene-modified KSCs to produce sufficient numbers of proliferated and differentiated keratinocytes to form a sheet. Low KSC colony formation and inefficient expansion of fresh isolated keratinocytes can unduly prolong epithelial sheet culture, resulting in the loss of stemness of KSCs and further limiting self-renewal potential. This could hinder the clinical application of epithelial sheet-based therapies. We thus investigated whether the *in-vitro* acceleration of keratinocyte culture using ROCKi could preserve KSCs’ self-renewal properties and their ability to undergo terminal differentiation.

Previous studies of ROCKi treatment in keratinocytes isolated from human neonatal foreskin observed increase of keratinocyte proliferation at the peak of day 6 with a 50-fold increase in CFE^23^. Studies have also shown that long-term exposure of keratinocytes to ROCKi increased proliferation without inducing any neoplastic transformation^39^. However, there was a report that longer treatments of ROCKi on cultured cells could potentially induce faster senescence after withdrawal of ROCKi treatment^40^. These studies suggested that ROCKi treatment is beneficial for effective expansion of keratinocytes, but long-term use of ROCKi in culture may exhaust KSCs’ proliferation potential, leading to senescence and loss of stemness. We thus used a short (six-day) ROCKi treatment to achieve increased cell survival and proliferation and hence efficiently expand KSCs *in-vitro* at the beginning of primary keratinocyte culture. We compared stem cells after a six-day treatment to untreated cells and also compared cells cultured for a further six days after ROCKi withdrawal with untreated cells. We confirmed that the influence of short-term ROCKi treatment on keratinocytes was transient and reversible.

We observed a 15-fold increase in CFE during ROCKi treatment. This was lower than a previously reported 50-fold increase^23^, possibly due to differences in tissue origin (the skin from donors with ear reconstruction surgery in this study versus foreskin in Terunuma’s study), culture conditions, and donor-related variability. Despite these differences, we confirmed by protein marker expression, single-cell transcriptomes and pseudotime analysis that the accelerated proliferation of primary keratinocytes following six-day ROCKi treatment was due to rapid KSC proliferation followed by differentiation into a large number of TA cells. Such an initial boost in keratinocyte growth can save at least 1–2 weeks in the generation of epithelial sheets.

We noted that ROCKi-treated keratinocytes were migrating non-cohesively with loose cell–cell contacts and mesenchymal-like cell morphology at the first three-day ROCKi treatment but became tightly packed at day 4. It has been reported that keratinocytes can undergo epithelial-to-mesenchymal transition (EMT) during wound healing and stay in this intermediate state (known as mesophase) to meet a high demand of cell proliferation required for wound closure^41^. It has also been reported that keratinocytes treated with ROCKi have fusiform-like phenotypes (spindle-shaped mesenchymal-like cells)^42^ which is similar to cells with depolymerized F-actin^43^. Rho-associated kinase (ROCK) can activate LIMK-kinase 1 (LIMK-1)^44^. As LIMK-1 inactivates the actin depolymerization factor ADF/cofilin binding to F-actin to control actin dynamics, inhibition of ROCK by ROCKi can promote activation and binding of ADF/Cofilin to F-actin, leading to F-actin depolymerization (i.e. loose cell–cell contacts / non-cohesive migration of cells). In addition, F-actin depolymerization has been shown to be involved in epithelial mesenchymal transition (EMT)^45^. EMT plays an essential role in wound healing where proliferation and migration of keratinocytes surrounding the wound is spatiotemporally controlled^46^. It has been shown that ROCKi increased adhesion and wound healing properties of human corneal endothelial cells^47^. We speculated that isolating and culturing primary keratinocytes *in-vitro* mimic the process of wound healing. In our primary keratinocyte culture, ROCKi may promote this ‘wound healing’ by stimulating keratinocytes into EMT in the first 2–3 days of treatment, later reverting to their epithelial state.

Mitochondrial content increases in stem cells changing from resting to activated states since resting stem cells need high energy for their transformation to proliferating progenitors^48-50^, which requires transition from glycolysis to oxidative phosphorylation (mitochondria-based ATP production). Generally, mitochondrial mass in the cell increases with the activation of mitochondrial biogenesis, a process of forming new and healthy mitochondria ^51,52^. We observed an increase in cell population with higher mitochondrial mass in ROCKi-treated culture, but we did not detect changes in the transcriptional level of genes related to mitochondrial biogenesis by scRNAseq. However, studies have found that the inhibition of mitophagy in cells could also increase mitochondrial mass^53^. This is because in somatic cells, mitophagy selectively eliminates damaged mitochondria, whereas in stem cells, mitophagy may serve as a physiologic mechanism for the metabolic rewiring of differentiating stem cells by removing old mitochondria and replacing them with new mitochondria configured for the differentiated cell state^49,52,54^. As in our study, cells with both higher mitochondrial mass and K14 (marker of proliferation) expression were localised in periphery of the colonies, indicating higher mitochondrial mass cells are proliferated cells, we suspect that the increased mitochondrial mass after short-term ROCKi treatment may be related to rapid differentiating KSCs to TA cells.

Transcriptome analysis at single-cell resolution revealed gene expression changes between treated cells and non-treated cells during ROCKi treatment. However, these differences disappeared following its withdrawal. This further supports that the influence of short-term ROCKi treatment is transient and reversible. We used *ANLN, AURKB, CCNA2, CKAP2L, FOXM1, HMGB2, LMNB1* as holoclone marker, *KRT14, TP63, ITGA6, ITB1, BIRC5* as meroclone markers, and *SERPINB3, SFN, KRT10, TGM1, IVL, SPINK5* as paraclone markers^37^ to classify cells as KSCs, TAs, and TDs and analysed the differentiation behaviour. This illustrated that the KSCs in ROCKi-treated cells differentiated to TA cells at a faster rate than non-treated cells. Combined with the evidence that increased holoclone colonies and proliferation-related marker changes in ROCKi-treated cells, it suggested that KSCs rapidly transformed to proliferating TA cells, thereby increasing the cell number and/or colony number. This agrees with the increased mitochondrial mass in ROCKi-treated cells. Importantly, the analysis further indicated that the proportion of KSCs in ROCKi-treated cultures was not reduced after ROCKi’s withdrawal, indicating that KSCs were not exhausted following short-term ROCKi application.

We used an *in-vitro* 3D organotypic system to check whether cells treated with ROCKi maintained the ability to terminally differentiate^55^. Both non-treated and ROCKi-treated cells were able to develop multiple layers of epidermis with the expressions of related protein markers. Notably, the proliferation marker Δ-P63α was clearly upregulated in the cultures generated using ROCKi-treated cells but reversed to the status similar to non-treated cells once ROCKi treatment was withdrawn. This further confirmed the transient effect of short-term ROCKi treatment. ROCK plays its biological function via AKT1/2 and ERK1/2 signalling, which is involved in cell survival, growth, and proliferation. We have shown, in primary keratinocytes, that ROCKi rapidly activated AKT1/2 and upstream kinases of ERK signalling, suggesting that the activation of AKT1/2 and ERK1/2 signalling was related to keratinocyte proliferation.

In conclusion, while short-term ROCKi treatment improved KSCs’ survival and growth, the influence on keratinocytes was transient, reversible, and did not affect KSCs’ characteristics. This could open a potential avenue for clinical application.

## Methods

### Skin samples and primary keratinocyte culture

Skin samples were obtained from healthy children (10-to 16-year-olds) undergoing plastic surgery for ear reconstruction, following informed consent. This study was approved by the local ethics committee with IRAS Project ID 279491. Fresh skin samples were processed within two hours for isolation and culture of keratinocytes as described by Di, et al. ^56^. Briefly, subcutaneous and adipose tissue were carefully removed from skin samples and the rest of the tissue was cut into small pieces (2 × 2 mm) and incubated with 0.02 U/mL neutral protease in PBS (Nordmark Pharma GmBH, Germany, #N0002936) for 3 hours at 37 °C to detach epidermis from the dermis. The epidermis was then transferred to 0.25% trypsin in 0.01% EDTA solution (Thermo Fisher Scientific, UK, #25200072) for 5 minutes to dissociate the epidermis into single cells. The dissociated cell solution was neutralized with Green’s medium^56^. Cells in the suspension were pelleted. After discarding the supernatant, pelleted cells were resuspended and plated out at a density of 4 × 10^4^/cm^2^ in culture flasks containing lethally irradiated 3T3-J2 cells (i3T3) and cultivated in a humidified atmosphere at 37°C with 10% CO_2_. Subconfluent cells were passaged and re-seeded at a density of 1.5 × 10^4^/cm^2^ with i3T3 in Green’s medium. Fresh isolated cells (P0) or cells with passage 1 (P1) were used in this study. All keratinocyte cultures were co-cultivated with i3T3 at 3 × 10^4^/cm^2^ unless specified.

### ROCK inhibitor Y-27632, treatment regimen and cell sampling

The ROCKi inhibitor Y-27632 ((R)-(+)-*trans*-4-(1-aminoethyl)-N-(4-pyridyl) cyclohexanecarboxamide-2) was purchased from AdooQ Bioscience (#A1101, CA). 10 mM stock solution was prepared by dissolving ROCKi powder in 100% dimethyl sulfoxide (DMSO, Merck Life Science, UK, #D8418) and filtered through 0.22µM syringe filters. The stock solution was then aliquoted and stored at - 20ºC for the use in cell culture. A 1:1000 dilution of the stock solution was used to make a final concentration of 10µM in culture medium.

Freshly isolated primary keratinocytes (P0) were seeded in 10cm dishes and cultured in the Green’s medium containing 10µM of ROCKi for six days. The culture medium was changed at day 3, replenishing with fresh ROCKi. Cells cultured in Green’s medium without ROCKi but containing 0.1% DMSO were used for controls and run with treated cells in parallel (**Figure 1**). Briefly, cells were seeded in triplicates for each group for assays including cell growth rate and morphology, mitochondrial mass, single cell RNA sequencing (scRNAseq), and immunoblotting. Six days after seeding, cultured cells from both ROCKi treated (ROCKi-6D^+^) and non-treated (Control-6D^−^) groups were harvested using 0.05% trypsin in DPBS. Before trypsinization, feeder layer was removed using 0.01% EDTA in DPBS for 30 seconds at room temperature. Trypsinized cells were neutralized with culture media and centrifuged at 500*g* for 5 min at 4°C. Cell pellets were resuspended in 0.5% fetal calf serum (FCS) in DPBS, and an aliquot was immediately proceeded for cell counting and cell size determination using the automated cell counter (CellDrop BF, DeNovix Inc, DE). Half of the cells were harvested at day 6 (ROCKi-6D^−^) and divided into three portions for mitochondrial mass assay, single-cell RNA sequencing analysis and immunoblotting. The remaining half cells were seeded back in triplicates at a density of 1×10^6^ cells per 10 cm dish and grown for a further six days without addition of ROCKi, and then were harvested for analysis. Cells without ROCKi (Control-6D^−^6D^−^ and ROCKi-6D^+^6D^−^) were processed in the same way.

### Immunoblotting

All antibodies used in this study are listed on **Table S1**.

Keratinocytes were lysed directly on the culture plate using 1x Laemmli sample buffer (BioRad, Herts, UK, #1610747) supplemented with fresh 50mM DTT at room temperature. Cell lysates were passed through 20 gauge needle several times and then heated at 96°C for 5 minutes. Total protein concentration in each sample was quantified using Pierce 660nM protein assay reagent (Thermo Fisher Scientific, Woolwich, UK, #22662) supplemented with ionic detergent compatibility reagent (IDCR, Thermo Fisher Scientific, Hampshire, UK, #22663). Equal amounts of total proteins were loaded in each lane, separated on Any kD Mini-PROTEAN TGX Stain-Free Protein Gels (BioRad, UK, #4568126) and transferred on to 0.2 µm Nitrocellulose membrane using Trans-Blot Turbo semi dry transfer system (BioRad, Herts UK). The transferred membranes were blocked for 1 hour with 5% w/v non-fat milk in tris-buffered saline containing Tween 20 (TBST, 19mM tris base, 137mM NaCl, 3mM KCl, 0.2% Tween 20, pH 7.4) for 1 hour and then incubated with primary antibodies diluted in blocking solution overnight at 4 °C. Membranes were washed three times (ten minutes each) in TBST before adding HRP-conjugated secondary antibodies for 2 hours at room temperature. Membranes were washed three times (ten minutes each) in TBST and target proteins were detected by using ECL Prime Western Blotting detection kit (SLS, UK, # RPN2232). Signal was visualized and recorded using ChemiDoc MP imaging system (BioRad, Herts, UK). The membranes were re-used by stripping the previous antibodies using Restore Plus western blot stripping buffer (Thermo Fisher Scientific, UK, #46430) and re-probed for the house-keeping protein glyceraldehyde 3-phosphate dehydrogenase (GAPDH) as a loading control. Protein bands were quantified against GAPDH by optical densitometry using Image Lab 6.1 software (BioRad, UK).

### MitoTracker Green staining

Primary keratinocytes were loaded with MitoTracker Green (200 nM) and incubated at 37 °C for 30 minutes. Cells were then washed with DPBS and feeder cells were removed using 0.01% EDTA in DPBS for 30 seconds at room temperature. Cells were then harvested by 0.05% of trypsin in DPBS, pelleted and then re-suspended in ice-cold 0.2% FCS in DPBS for flow cytometry analysis using CytoFLEX S (Beckman Coulter, Bucks, UK). Cells were excited at 488 nm and the emission fluorescence was collected at 535 nm. 40,000 live cells were analysed per sample.

### 3D organotypic culture

A 3D organotypic culture was performed as described by Di, et al. ^55^. A de-epidermalised dermis (DED, Euro Tissue Bank, BEVERWIJK, Netherlands) was used as a scaffold for generating 3D organotypic culture. A metal ring with 1cm diameter was placed on the dermal side of the DED in a 35mm dish and 2 × 10^5^ primary human dermal fibroblast were seeded on the reticular side and cultured in the Green’s medium. After 24 hours, the metal ring was removed, DED was flipped over (in the papillary side), and the metal ring was placed again on the papillary side. 5 ×10^5^ primary keratinocytes were then seeded on the papillary side of the DED in Green’s medium. After 48 hours, the metal ring was removed, and the DED culture was air-lifted by transferring it on to a mesh so that reticular side of the DED was in contact with the culture medium and the papillary side was in contact with air. Air-lifted DED with cells were cultured for 14 days, allowing the keratinocytes to proliferate and differentiate. Primary keratinocytes from three different donors were used to generate three biological replicates. Primary human fibroblast (passage 3 to 4) from same donor was used for all 3D cultures.

### Cryosectioning and immunostaining

In supplementary materials.

### Colony-formation assay

The colony forming efficiency (CFE) of the keratinocytes was determined by a colony-formation assay as previously described^11^. Keratinocytes were seeded in 10cm dishes at a density of 800 cells per dish. On day 6 and day 12, dishes cultured with cells were washed twice with DPBS and stained with 0.25% rhodanile blue (Merck Life Science, UK, #121495) in 1% sulphuric acid/DPBS solution for 20 minutes at room temperature. Dishes were washed several times with DPBS and air dried. Colony staining image were scanned at the setting of 600 DPI using an HP Envy 4500 (HP, UK) scanner. A ruler was scanned at 600 DPI to measure the number of pixels per millimetre. Colony numbers and areas were counted and measured using Fiji software (https://imagej.net/software/fiji/). Briefly, images were converted to 8-bit grayscale. Uneven brightness of the staining was adjust using adaptive median filter (r=2), the threshold was set to default, hollow colonies were filled by using binary fill holes, and joined colonies were separated using binary watershed. Colonies ≥ 1mm^2^ in area with the circularity of ≥ 0.5 were counted as colonies.

To determine clonality of the colonies, six-day old colonies were trypsinised and seeded back in indicator dishes and cultured for additional 12 days. Based on the size and circularity of the colonies on indicator dishes, the parent colonies were identified as holoclone, meroclone, or paraclone^11^. In this study, we did not perform colony-type identification as we did the CFE assay on day 6 after ROCKi treatment.

### Single-cell library preparation and RNA sequencing

Cultured keratinocytes were harvested and resuspended in ice-cold buffer containing 0.5% FCS in DPBS. A small aliquot of the cell suspension was used to determine the percentage of live/dead cells by stained with Acridine Orange/Propidium Iodide (AO/PI) Cell Viability Kit (Vita Scientific, MD, #LGBD10012) and counted by using LUNA-FL automated cell counter (Logos Biosystems, VA). Approximately 3,000 live cells from each group were loaded into one channel of the Chromium Chip G using the Single Cell reagent kit v3.1 (10X Genomics, CA, US) for capturing single cell Gel-Bead Emulsion using the Chromium controller. Following single cell capture, cells were lysed, and cDNA of individual cells were synthesized within the Gel-Bead Emulsion and was used for library preparation. cDNA was amplified and size-selected using SPRISelect (Beckman Coulter, Germany, #B23317) following the manufacturer’s protocol. 25ng of the size-selected cDNA of each sample were used to construct Illumina sequencing libraries. Libraries were pooled at equal concentration and sequenced on the NovaSeq SP100 or NextSeq2000 Illumina sequencing platform at 50,000 reads per cell.

### Generation of gene expression matrix of counts and quality control for scRNAseq

The gene expression count matrix was generated from scRNAseq raw sequences using cellranger^57^ with the standard cellranger pipeline consisting of the *mkfastq, count*, and *aggr* steps. Gene expression was analysed using Seurat^58^. Genes present in fewer than three cells and cells expressing fewer than 200 genes were excluded from the analysis. Low-quality cells and barcodes corresponding to multiple cells or empty droplets were removed by excluding cells associated with either an anomalous number of genes, number of molecules, or percentage of mitochondrial DNA. Making the Gel-Bead Emulsion for each sample in separate wells made these measures vary between each other. Hence, we used different thresholds for each sample as a universal threshold would include low-quality cells in some samples and exclude high-quality cells in others (**Supplementary Figure S4**). Non-keratinocytes were removed by excluding cells expressing (>1 count) of markers of melanocytes (*MLANA, PMEL, MITF*) and mesenchymal-like cells (*ACTG2, DLK1, MTRNR2L6, MTRNR2L10, MTRNR2L7, MTRNR2L1, PRKAR2B, NR2F1*)^59^.

### Normalisation and cell cycle removal

Gene expression was normalised using sctransform^60^. The effect of cell cycle was mitigated by assigning scores to each cell representing the probability of its being in different stages of the cell cycle (using Seurat’s *CellCycleScoring* function) and then regressing out the difference between these scores using sctransform (satijalab.org/seurat/archive/v3.1/cell_cycle_vignette.html - “Alternative Workflow”).

### Dimensionality reduction and donor effect removal

Dimensionality reduction was carried out by principal components analysis (PCA) using the 3,000 most highly variable genes as features. The 50 most significant components were used since there was no clear point at which the generated components became less significant (**Supplementary Figure S5**). Inter-donor variation was removed with Harmony^61^.

### Differential expression and differential abundance analyses

Seurat was used for differential expression analysis of treated and control cells with its standard Wilcoxon rank sum test. We used DAseq^30^ to measure the proportion of two cell populations that cluster separately (detailed in supplementary methods).

### Cell type proportion comparison

SCINA^62^ was used to compare the cell type proportions in the control and ROCKi-treated cells. Cell type markers published by Enzo, et al. ^37^ were used to classify cells as either holoclone-forming (HF), mero-or paraclone-forming (MP), or terminally differentiated (TD). Bootstrap resampling was used to estimate confidence intervals around the total absolute differences between cell type proportions.

### Trajectory analysis

Slingshot^63^ was used to infer the differentiation trajectory for each sample class with the Harmony reduction. The groups of non-treated and ROCKi-treated cells were combined for the inference procedure to ensure that the pseudotimes were directly comparable. Three clusters were used for each sample (approximately corresponding to holoclone-forming, meroclone-forming, and paraclone-forming cells) as the input to Slingshot. The cluster likely to contain mostly holoclone-forming cells were used as the starting point for each inferred trajectory. Trajectories were then compared by computing the 0.1^th^, 0.2^th^, …, 99.9^th^, 100^th^ percentiles of the pseudotimes corresponding to each sample and calculating the mean absolute difference at each percentile. Bootstrap resampling was used to estimate confidence intervals around the mean absolute differences between the pseudotimes both during ROCKi treatment and following its withdrawal.

## Data availability

The rmarkdown notebook used to generate our single-cell RNAseq results is available at github.com/george-hall-ucl/reversibility_of_rocki_paper_scrnaseq/blob/main/data_analysis_for_paper.md.

## Acknowledgements

Supported by MRC (grant ref: MR/S036989/1) and the National Institute for Health Research (NIHR) Biomedical Research Centre based at Great Ormond Street Hospital/Institute of Child Health. The views expressed are those of the author(s) and not necessarily those of the NHS, the NIHR or the Department of Health. VJ is supported by MRC and GTH is supported by the Biomedical Research Centre at Great Ormond Street Hospital.

## Author Contributions

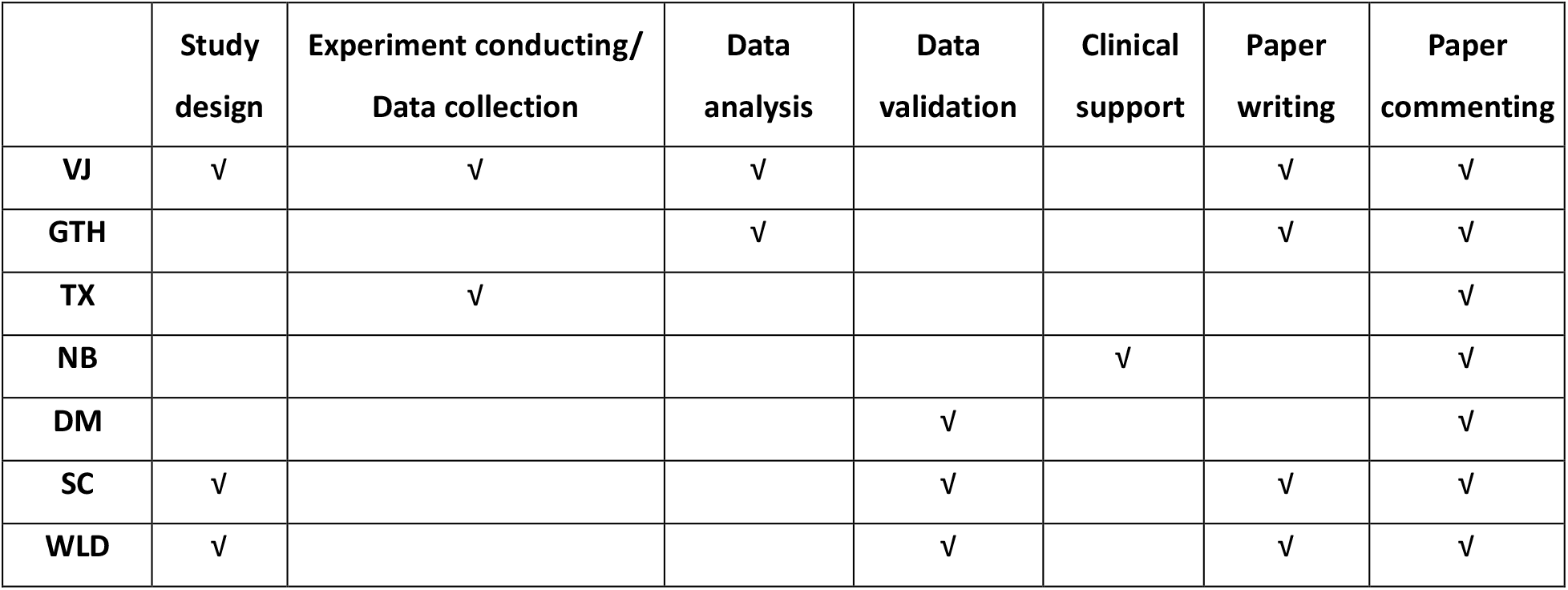

## Competing Interests statement

We have no conflicts of interest to declare.

## Notes

### Competing Interest Statement

The authors have declared no competing interest.

## References

1 Lo, C. H., Chong, E., Akbarzadeh, S., Brown, W. A. & Cleland, H. A systematic review: Current trends and take rates of cultured epithelial autografts in the treatment of patients with burn injuries. Wound Repair Regen 27, 693–701, doi:10.1111/wrr.12748 (2019).

2 Hirsch, T. et al. Regeneration of the entire human epidermis using transgenic stem cells. Nature 551, 327–332, doi:10.1038/nature24487 (2017).

3 Siprashvili, Z. et al. Safety and Wound Outcomes Following Genetically Corrected Autologous Epidermal Grafts in Patients With Recessive Dystrophic Epidermolysis Bullosa. Jama 316, 1808–1817, doi:10.1001/jama.2016.15588 (2016).

4 Di, W.-L. et al. Generation and Clinical Application of Gene-Modified Autologous Epidermal Sheets in Netherton Syndrome: Lessons Learned from a Phase 1 Trial. Human Gene Therapy 30, 1067–1078, doi:10.1089/hum.2019.049 (2019).

5 Jayarajan, V., Kounatidou, E., Qasim, W. & Di, W.-L. Ex vivo gene modification therapy for genetic skin diseases—recent advances in gene modification technologies and delivery. Experimental Dermatology 30, 887–896, doi:https://doi.org/10.1111/exd.14314 (2021).

6 Li, A., Simmons, P. J. & Kaur, P. Identification and isolation of candidate human keratinocyte stem cells based on cell surface phenotype. Proceedings of the National Academy of Sciences of the United States of America 95, 3902–3907, doi:10.1073/pnas.95.7.3902 (1998).

7 De Rosa, L. et al. Toward Combined Cell and Gene Therapy for Genodermatoses. Cold Spring Harbor perspectives in biology 12, doi:10.1101/cshperspect.a035667 (2020).

8 Frisch, S. M. & Francis, H. Disruption of epithelial cell-matrix interactions induces apoptosis. The Journal of cell biology 124, 619–626, doi:10.1083/jcb.124.4.619 (1994).

9 Watt, F. M. Stem cell fate and patterning in mammalian epidermis. Current Opinion in Genetics & Development 11, 410–417, doi:https://doi.org/10.1016/S0959-437X(00)00211-2 (2001).

10 Rheinwald, J. G. & Green, H. Serial cultivation of strains of human epidermal keratinocytes: the formation of keratinizing colonies from single cells. Cell 6, 331–343, doi:10.1016/s0092-8674(75)80001-8 (1975).

11 Barrandon, Y. & Green, H. Three clonal types of keratinocyte with different capacities for multiplication. Proc Natl Acad Sci U S A 84, 2302–2306, doi:10.1073/pnas.84.8.2302 (1987).

12 O’Connor, N., Mulliken, J., Banks-Schlegel, S., Kehinde, O. & Green, H. GRAFTING OF BURNS WITH CULTURED EPITHELIUM PREPARED FROM AUTOLOGOUS EPIDERMAL CELLS. The Lancet 317, 75–78, doi:https://doi.org/10.1016/S0140-6736(81)90006-4 (1981).

13 Watanabe, K. et al. A ROCK inhibitor permits survival of dissociated human embryonic stem cells. Nat Biotechnol 25, 681–686, doi:10.1038/nbt1310 (2007).

14 Koyanagi, M. et al. Inhibition of the Rho/ROCK pathway reduces apoptosis during transplantation of embryonic stem cell-derived neural precursors. Journal of Neuroscience Research 86, 270–280, doi:https://doi.org/10.1002/jnr.21502 (2008).

15 Ohgushi, M. et al. Molecular pathway and cell state responsible for dissociation-induced apoptosis in human pluripotent stem cells. Cell Stem Cell 7, 225–239, doi:10.1016/j.stem.2010.06.018 (2010).

16 Zhang, L. et al. ROCK inhibitor Y-27632 suppresses dissociation-induced apoptosis of murine prostate stem/progenitor cells and increases their cloning efficiency. PLoS One 6, e18271, doi:10.1371/journal.pone.0018271 (2011).

17 Wu, Y. et al. ROCK inhibitor Y27632 promotes proliferation and diminishes apoptosis of marmoset induced pluripotent stem cells by suppressing expression and activity of caspase 3. Theriogenology 85, 302–314, doi:10.1016/j.theriogenology.2015.09.020 (2016).

18 Yu, Z. et al. ROCK inhibition with Y27632 promotes the proliferation and cell cycle progression of cultured astrocyte from spinal cord. Neurochem Int 61, 1114–1120, doi:10.1016/j.neuint.2012.08.003 (2012).

19 Wen, L. et al. Establishment of an Efficient Primary Culture System for Human Hair Follicle Stem Cells Using the Rho-Associated Protein Kinase Inhibitor Y-27632. Front Cell Dev Biol 9, 632882, doi:10.3389/fcell.2021.632882 (2021).

20 Sun, C. C., Chiu, H. T., Lin, Y. F., Lee, K. Y. & Pang, J. H. Y-27632, a ROCK Inhibitor, Promoted Limbal Epithelial Cell Proliferation and Corneal Wound Healing. PLoS One 10, e0144571, doi:10.1371/journal.pone.0144571 (2015).

21 Nemati, S. et al. Long-term self-renewable feeder-free human induced pluripotent stem cell-derived neural progenitors. Stem Cells Dev 20, 503–514, doi:10.1089/scd.2010.0143 (2011).

22 o, E. et al. The Rho kinase inhibitor fasudil augments the number of functional endothelial progenitor cells in ex vivo cultures. Int J Mol Med 28, 357–363, doi:10.3892/ijmm.2011.698 (2011).

23 Terunuma, A., Limgala, R. P., Park, C. J., Choudhary, I. & Vogel, J. C. Efficient procurement of epithelial stem cells from human tissue specimens using a Rho-associated protein kinase inhibitor Y-27632. Tissue Eng Part A 16, 1363–1368, doi:10.1089/ten.TEA.2009.0339 (2010).

24 Nagata, K. et al. Effects of fasudil hydrochloride on cerebral blood flow in patients with chronic cerebral infarction. Clin Neuropharmacol 16, 501–510, doi:10.1097/00002826-199312000-00003 (1993).

25 Ahmadieh, H. et al. Intravitreal injection of a Rho-kinase inhibitor (fasudil) combined with bevacizumab versus bevacizumab monotherapy for diabetic macular oedema: a pilot randomised clinical trial. Br J Ophthalmol 103, 922–927, doi:10.1136/bjophthalmol-2018-312244 (2019).

26 Olson, M. F. Applications for ROCK kinase inhibition. Curr Opin Cell Biol 20, 242–248, doi:10.1016/j.ceb.2008.01.002 (2008).

27 Suzuki, Y., Shibuya, M., Satoh, S.-i., Sugimoto, Y. & Takakura, K. A postmarketing surveillance study of fasudil treatment after aneurysmal subarachnoid hemorrhage. Surgical Neurology 68, 126–131, doi:https://doi.org/10.1016/j.surneu.2006.10.037 (2007).

28 Amano, M., Nakayama, M. & Kaibuchi, K. Rho-kinase/ROCK: A key regulator of the cytoskeleton and cell polarity. Cytoskeleton (Hoboken) 67, 545–554, doi:10.1002/cm.20472 (2010).

29 Pellegrini, G. et al. p63 identifies keratinocyte stem cells. Proc Natl Acad Sci U S A 98, 3156–3161, doi:10.1073/pnas.061032098 (2001).

30 Jun Zhao, A. J., Henry Li, Ofir Lindenbaum, Esen Sefik, Ruaidhrí Jackson, Xiuyuan Cheng, Richard A. Flavell, and Yuval Kluger. Detection of differentially abundant cell subpopulations in scRNA-seq data. Proceedings of the National Academy of Sciences 118 (2021).

31 Satelli, A. & Li, S. Vimentin in cancer and its potential as a molecular target for cancer therapy. Cell Mol Life Sci 68, 3033–3046, doi:10.1007/s00018-011-0735-1 (2011).

32 Sharma, P. et al. Keratin 19 regulates cell cycle pathway and sensitivity of breast cancer cells to CDK inhibitors. Scientific Reports 9, 14650, doi:10.1038/s41598-019-51195-9 (2019).

33 Huang, B. et al. Long non-coding antisense RNA KRT7-AS is activated in gastric cancers and supports cancer cell progression by increasing KRT7 expression. Oncogene 35, 4927–4936, doi:10.1038/onc.2016.25 (2016).

34 Post, A., Pannekoek, W. J., Ponsioen, B., Vliem, M. J. & Bos, J. L. Rap1 Spatially Controls ArhGAP29 To Inhibit Rho Signaling during Endothelial Barrier Regulation. Mol Cell Biol 35, 2495–2502, doi:10.1128/mcb.01453-14 (2015).

35 Abhishek, S. & Palamadai Krishnan, S. Epidermal Differentiation Complex: A Review on Its Epigenetic Regulation and Potential Drug Targets. Cell J 18, 1–6, doi:10.22074/cellj.2016.3980 (2016).

36 Ogawa, E. et al. Epidermal FABP (FABP5) Regulates Keratinocyte Differentiation by 13(S)-HODE-Mediated Activation of the NF-κB Signaling Pathway. Journal of Investigative Dermatology 131, 604–612, doi:https://doi.org/10.1038/jid.2010.342 (2011).

37 Enzo, E. et al. Single-keratinocyte transcriptomic analyses identify different clonal types and proliferative potential mediated by FOXM1 in human epidermal stem cells. Nature Communications 12, 2505, doi:10.1038/s41467-021-22779-9 (2021).

38 Totsukawa, G. et al. Distinct roles of ROCK (Rho-kinase) and MLCK in spatial regulation of MLC phosphorylation for assembly of stress fibers and focal adhesions in 3T3 fibroblasts. The Journal of cell biology 150, 797–806, doi:10.1083/jcb.150.4.797 (2000).

39 Chapman, S., Liu, X., Meyers, C., Schlegel, R. & McBride, A. A. Human keratinocytes are efficiently immortalized by a Rho kinase inhibitor. J Clin Invest 120, 2619–2626, doi:10.1172/jci42297 (2010).

40 Chapman, S., McDermott, D. H., Shen, K., Jang, M. K. & McBride, A. A. The effect of Rho kinase inhibition on long-term keratinocyte proliferation is rapid and conditional. Stem Cell Res Ther 5, 60–60, doi:10.1186/scrt449 (2014).

41 Nakamura, M. & Tokura, Y. Epithelial–mesenchymal transition in the skin. Journal of Dermatological Science 61, 7–13, doi:https://doi.org/10.1016/j.jdermsci.2010.11.015 (2011).

42 Gandham, V. D. et al. Effects of Y27632 on keratinocyte procurement and wound healing. Clin Exp Dermatol 38, 782–786, doi:10.1111/ced.12067 (2013).

43 Hopkins, A. M. et al. Organized migration of epithelial cells requires control of adhesion and protrusion through Rho kinase effectors. American Journal of Physiology-Gastrointestinal and Liver Physiology 292, G806–G817, doi:10.1152/ajpgi.00333.2006 (2007).

44 Ohashi, K. et al. Rho-associated kinase ROCK activates LIM-kinase 1 by phosphorylation at threonine 508 within the activation loop. J Biol Chem 275, 3577–3582, doi:10.1074/jbc.275.5.3577 (2000).

45 Haynes, J., Srivastava, J., Madson, N., Wittmann, T. & Barber, D. L. Dynamic actin remodeling during epithelial-mesenchymal transition depends on increased moesin expression. Mol Biol Cell 22, 4750–4764, doi:10.1091/mbc.E11-02-0119 (2011).

46 Haensel, D. & Dai, X. Epithelial-to-mesenchymal transition in cutaneous wound healing: Where we are and where we are heading. Developmental Dynamics 247, 473–480, doi:https://doi.org/10.1002/dvdy.24561 (2018).

47 Pipparelli, A. et al. ROCK Inhibitor Enhances Adhesion and Wound Healing of Human Corneal Endothelial Cells. PLOS ONE 8, e62095, doi:10.1371/journal.pone.0062095 (2013).

48 Hinge, A. et al. Asymmetrically Segregated Mitochondria Provide Cellular Memory of Hematopoietic Stem Cell Replicative History and Drive HSC Attrition. Cell Stem Cell 26, 420-430.e426, doi:10.1016/j.stem.2020.01.016 (2020).

49 Mandal, S., Lindgren, A. G., Srivastava, A. S., Clark, A. T. & Banerjee, U. Mitochondrial function controls proliferation and early differentiation potential of embryonic stem cells. Stem Cells 29, 486–495, doi:10.1002/stem.590 (2011).

50 Wang, Z. A., Huang, J. & Kalderon, D. Drosophila follicle stem cells are regulated by proliferation and niche adhesion as well as mitochondria and ROS. Nat Commun 3, 769, doi:10.1038/ncomms1765 (2012).

51 Ploumi, C., Daskalaki, I. & Tavernarakis, N. Mitochondrial biogenesis and clearance: a balancing act. The FEBS Journal 284, 183–195, doi:https://doi.org/10.1111/febs.13820 (2017).

52 Bahat, A. & Gross, A. Mitochondrial plasticity in cell fate regulation. J Biol Chem 294, 13852–13863, doi:10.1074/jbc.REV118.000828 (2019).

53 Townley, A. R. & Wheatley, S. P. Mitochondrial survivin reduces oxidative phosphorylation in cancer cells by inhibiting mitophagy. J Cell Sci 133, doi:10.1242/jcs.247379 (2020).

54 Krantz, S. et al. Mitophagy mediates metabolic reprogramming of induced pluripotent stem cells undergoing endothelial differentiation. Journal of Biological Chemistry 297, doi:10.1016/j.jbc.2021.101410 (2021).

55 Di, W. L. et al. Ex-vivo gene therapy restores LEKTI activity and corrects the architecture of Netherton syndrome-derived skin grafts. Molecular therapy : the journal of the American Society of Gene Therapy 19, 408–416, doi:10.1038/mt.2010.201 (2011).

56 Di, W. L. et al. Phase I study protocol for ex vivo lentiviral gene therapy for the inherited skin disease, Netherton syndrome. Hum Gene Ther Clin Dev 24, 182–190, doi:10.1089/humc.2013.195 (2013).

57 Zheng, G. X. Y. et al. Massively parallel digital transcriptional profiling of single cells. Nature Communications 8, 14049, doi:10.1038/ncomms14049 (2017).

58 Hao, Y. et al. Integrated analysis of multimodal single-cell data. Cell 184, 3573-3587.e3529, doi:https://doi.org/10.1016/j.cell.2021.04.048 (2021).

59 Karlsson, M. et al. A single-cell type transcriptomics map of human tissues. Sci Adv 7, doi:10.1126/sciadv.abh2169 (2021).

60 Hafemeister, C. & Satija, R. Normalization and variance stabilization of single-cell RNA-seq data using regularized negative binomial regression. Genome Biology 20, 296, doi:10.1186/s13059-019-1874-1 (2019).

61 Korsunsky, I. et al. Fast, sensitive and accurate integration of single-cell data with Harmony. Nature Methods 16, 1289–1296, doi:10.1038/s41592-019-0619-0 (2019).

62 Zhang, Z. et al. SCINA: A Semi-Supervised Subtyping Algorithm of Single Cells and Bulk Samples. Genes (Basel) 10, doi:10.3390/genes10070531 (2019).

63 Street, K. et al. Slingshot: cell lineage and pseudotime inference for single-cell transcriptomics. BMC Genomics 19, 477, doi:10.1186/s12864-018-4772-0 (2018).

