## Supplementary materials for "Short-term Rho-associated kinase inhibitor treatment accelerates primary keratinocyte growth while preserving stem cell characteristics"

### Materials and Methods

#### Cryosectioning, formalin-fixed paraffin embedding and sectioning

Organotypic cultures at day 14 after airlifting were harvested, snap frozen, embedded in optimal cutting temperature compound (Tissue-Tek, Belgium, #94-4583) and stored at -70°C for sectioning. Frozen tissues were cut into 10µm-thick slices at -20°C using Leica CM1850 Cryostat (Leica Instruments, UK) and transferred onto polylysine adhesion slides (Thermo Fisher Scientific, UK, # 10219280). Slides were stored at -20°C for immunostaining.

#### H&E and Immunofluorescence staining

Frozen sections were air-dried for 30 minutes. Following rinsing with water, haematoxylin (Gill no. 3, Sigma, UK, #GHS332) was added on the sections for 40 minutes. Slides were then rinsed with water and dipped in 2% HCL in 70% ethanol once to induce haematoxylin differentiation. Slides were further rinsed with hot tap water, immersed in Scott's tap water for 1 minute and rinsed with hot tap water again before incubating with 1% eosin (Sigma, UK, #HT110232) for 2 minutes. Slides were then dehydrated by rinsing with water, 70%, and 100% ethanol for 30 seconds each. Dehydrated slides were immersed in xylene for 10 minutes and mounted using DPX new (Merck Life Science, UK, #100579). Frozen sections were air-dried at room temperature for 30 minutes and rinsed with PBS for 5 minutes. Sections were blocked with 3% FCS in PBS and incubated with the primary antibody (**Table S1**) in 1% bovine serum albumin overnight at 4°C. The following day, slides were washed with PBS three times (for five minutes each time) and incubated with fluorescent-conjugated secondary antibody (**Table S1**) for 1 hour at room temperature. Slides were then washed three times (for five minutes each time) and counterstained with Hoechst 33342 in 1:10,000 dilution in PBS for 5 minutes and washed three times (for five minutes each time). Negative controls which were without primary antibodies were processed in parallel. Sections were mounted using Fluoromount-G mounting medium (Thermo Fisher Scientific, Woolwich, UK, # 00-4958-02) and examined using ZEISS LSM710 inverted confocal microscope (Carl Zeiss Ltd, UK).

**Gene expression matrix generation**

For each control and ROCKi-treated sample, we generated a gene expression count matrix with the standard Cellranger (v5.0.1) pipeline[1]. We converted BCL files into fastq files using Cellranger *mkfastq* with default parameter values and extracted gene expression data for each sample using Cellranger *count* with the *refdata-cellranger-GRCh38-3.0.0 reference* transcriptome with default parameter values. We aggregated the counts across samples using Cellranger *aggr* with no normalisation (i.e. *--normalize=none*), and otherwise used default parameter values.

**Normalisation and cell cycle regression**

We normalised counts using sctransform, employing the sctransform package (v0.3.2) and the *SCTransform* function with default parameter settings. We then mitigated the effect of cell cycle by assigning scores to each cell representing the likelihood of it being in different stages of the cell cycle (using Seurat's *CellCycleScoring* function with default parameter values and its default cell cycle markers "s.genes" and "g2m.genes") and regressing out the difference between these scores, again using sctransform with its default parameter values. The developers of Seurat recommend this approach when dealing with cells undergoing differentiation as it does not remove the difference between cycling and non-cycling cells while still reducing the impact of the cell cycle phase in proliferating cells (*https://satijalab.org/seurat/ archive/v3.1/cell_cycle_vignette.html* - "Alternative Workflow"). For the remainder of the analysis, we used the resulting normalised assay ("SCT") instead of the un-normalised RNA counts.

**Dimensionality reduction**

We carried out dimensionality reduction using principal components analysis with Seurat's *RunPCA* function using the 3,000 most highly variable genes (from sctransform) as features. We used the fifty most significant components (i.e. those that explain the most variance of the dataset) since there was no clear point at which the generated components became less important (**Supplementary Figure S5**).

**Donor effect removal and visualisation**

We reduced the inter-donor variation with the Harmony package (v0.1.0)[2], using *RunHarmony* with the SCT assay and labels corresponding to donor of origin. We used the projection produced by Harmony when dealing with cells from multiple donors.

We visualised the cells in a UMAP plot, generated by Seurat's *RunUMAP* function with default parameter values.

**Differential expression and differential abundance analysis**

We carried out differential expression analysis using the standard Wilcoxon rank sum test in Seurat's *FindMarkers* function with default parameter values.

Differential abundance was estimated using DAseq (v1.0.0)[3]. We used the Harmony reduction and default parameter values.

### Proportions of cell types in control and ROCKi-treated cells

We used the SCINA package (v1.2.0)[4] to estimate the cell type proportions in the control/ROCKi-treated cells (differential composition analysis[5]). We used cell type markers identified by Enzo *et al.* [6] to classify cells as either holoclone-forming (HF), Mero- or Paraclone-forming (MP), or terminally differentiated (TD). We used the *SCINA* function with default parameter values.

We quantified the difference between the cell type proportions in the treated and control cells at both day 6 and day 12. Formally, denoting the proportion of HF cells in day 6 control and ROCKi-treated cells in the sample as $p_{\mathrm{HF}}^{6c}$ and $p_{\mathrm{HF}}^{6t}$, respectively, and denoting the corresponding proportions of MP cells in the sample as $p_{MP}^{6c}$ and $p_{MP}^{6t}$, we are interested in $D_6 := |p_{\mathrm{HF}}^{6c} - p_{\mathrm{HF}}^{6t}| + |p_{\mathrm{MP}}^{6c} - p_{\mathrm{MP}}^{6t}|$ (and the corresponding day 12 difference $D_{12}$, defined similarly). We do not include the difference in the proportions of TD cells since this is implied by the previous two differences and thus additionally estimating this proportion is only liable to increase the error in our confidence intervals.

We resampled cell type labels with replacement and ensured that the proportion of control and ROCKi-treated cells in both populations was preserved. We repeated this process 10,000 times. Denoting the proportion of HF cells in Control-6D⁻in the $i^{\mathrm{th}}$ resample as $\hat{p}_{\mathrm{HF,i}}^{6c}$, and defining $\hat{\mathrm{p}}_{\mathrm{HF,i}}^{6\mathrm{t}}$, $\hat{p}_{MP,i}^{6c}$, and $\hat{p}_{MP,i}^{6t}$ similarly, we define the total absolute difference in cell type proportions of the $i^{\mathrm{th}}$ resample of $D_6$ as $\widehat{D}_6^i := |\hat{p}_{HF,i}^{6c} - \hat{p}_{HF,i}^{6t}| + |\hat{p}_{MP,i}^{6c} - \hat{p}_{MP,i}^{6t}|$, with corresponding residual $R_6^i := \widehat{D}_6^i - D_6$. We define the set of all residuals as $R_6 := \{R_6^i \,|1 \leq i \leq 10{,}000\}$ and denote the 2.5th and 97.5th percentiles as $\delta_{2.5}^{R_6}$ and $\delta_{97.5}^{R_6}$, respectively. Thus the 95% confidence interval for the total absolute difference in the proportions of HF and MP cells in treated and control cells in the whole cell population as $[D_6 - \delta_{97.5}^{R_6}, D_6 - \delta_{2.5}^{R_6}]$. The confidence interval for this quantity at day 12 was computed in the same way.

**Trajectory inference**

We used Slingshot (v1.8.0)[7] to infer the differentiation trajectory for day 6 and day 12 cells, using the Harmony reduction. We combined the control and ROCKi-treated cells for the inference procedure to ensure that the pseudotimes were directly comparable. If trajectories were inferred separately, then the pseudotimes of both sets of cells would need to be normalised to allow comparison, which could lead to cells at different stages of the differentiation process (e.g. HF cells and TD cells) incorrectly appearing to have similar pseudotimes. In addition, the trajectories inferred by this approach are likely to be more reliable since they are based on a larger number of cells. This is the recommended approach in other differential progression methods such as condiments[8]. We used the Seurat *FindClusters* function with a resolution of 0.1 (and otherwise default parameter values) to produce three clusters for each sample class (which can be thought of as corresponding to HF, MP, and TD cells, respectively) and used these clusters as the input to Slingshot. We used the *Slingshot* function from the slingshot package with default parameter values and specified the cluster judged manually to comprise mainly HF cells as the starting point for each inferred trajectory.

**Trajectory comparison**

We visually compared the trajectories of control and ROCKi-treated cells at both day 6 and day 12 by plotting the 0.1$^{th}$, 0.2$^{th}$, …, 99.9$^{th}$, 100$^{th}$ percentiles of the pseudotimes corresponding to each sample class. The resulting curves correspond to approximations of the empirical cumulative distribution functions of the pseudo-times with flipped axes (since the percentage through the differentiation process is on the x-axis, and pseudo-time is on the y-axis). We found this to be a more intuitive method of comparing trajectories than the kernel density estimation of the pseudotimes as used in the "condiments" package[9].

We quantified the difference between the differentiation trajectories of the control and ROCKi-treated cells at both day 6 and day 12 using the mean absolute difference (MAD) between the pseudotimes at each percentile as an approximation to the area between the curves.

We used bootstrap resampling to compute confidence intervals of this quantity, both at day 6 and day 12. Formally, let $p_{6c}^{i}$ be pseudotimes corresponding to the $(i/10)$-th percentile of the pseudotimes of Control-6D$^{-}$ and define $p_{6t}^{i}$ similarly for ROCKi-6D$^{+}$. Then the MAD of the pseudotimes is $\sum_{i=1}^{1000} \left| p_{6c}^{i} - p_{6t}^{i} \right| / 1000$. A MAD of 0 would indicate that the two differentiation trajectories are identical, whilst larger MADs indicate more dissimilar trajectories.

We used bootstrap resampling to derive confidence intervals on the MAD of the pseudotimes of treated and untreated cells, at both day 6 and day 12. To do so, we resampled day 6 cells with replacement and made sure to maintain the proportions of control and ROCKi-treated cells. We then computed the pseudotime of each cell using the method described above and computed the MAD of the two sets of pseudotimes. After computing 10,000 MADs, we calculated the residuals with respect to the MAD from the observed data, denoted $\mathrm{M}$. Denoting the 2.5% and 97.5% percentiles of the residuals as $\delta_{2.5}$ and $\delta_{97.5}$, respectively, we compute the 95% confidence interval of the MAD as $[\mathrm{M} - \delta_{97.5}, \mathrm{M} - \delta_{2.5}]$. We compared the differentiation trajectories of the day 12 cells in the same way.

### References


1 Zheng, G. X. Y. *et al.* Massively parallel digital transcriptional profiling of single cells. *Nature Communications* **8**, 14049, doi:10.1038/ncomms14049 (2017).

2 Ilya Korsunsky, N. M., Jean Fan, Kamil Slowikowski, Fan Zhang, Kevin Wei, Yuriy Baglaenko, Michael Brenner, Po-ru Loh & Soumya Raychaudhuri. Fast, sensitive and accurate integration of single-cell data with Harmony. *Nature Methods* **16**, 1289–1296 (2019).

3 Jun Zhao, A. J., Henry Li, Ofir Lindenbaum, Esen Sefik, Ruaidhrí Jackson, Xiuyuan Cheng, Richard A. Flavell, and Yuval Kluger. Detection of differentially abundant cell subpopulations in scRNA-seq data. *Proceedings of the National Academy of Sciences* **118** (2021).

4 Zhang, Z. *et al.* SCINA: A Semi-Supervised Subtyping Algorithm of Single Cells and Bulk Samples. *Genes (Basel)* **10**, doi:10.3390/genes10070531 (2019).

5 Yue Cao, Y. L., John T. Ormerod, Pengyi Yang, Jean Y.H. Yang & Kitty K. Lo. scDC: single cell differential composition analysis. *BMC Bioinformatics volume* (2019).

6 Enzo, E. *et al.* Single-keratinocyte transcriptomic analyses identify different clonal types and proliferative potential mediated by FOXM1 in human epidermal stem cells. *Nature Communications* **12**, 2505, doi:10.1038/s41467-021-22779-9 (2021).

7 Street, K. *et al.* Slingshot: cell lineage and pseudotime inference for single-cell transcriptomics. *BMC Genomics* **19**, 477, doi:10.1186/s12864-018-4772-0 (2018).

8 Hector Roux de Bézieux, K. V. d. B., Kelly Street, Sandrine Dudoit. Trajectory inference across multiple conditions with condiments: differential topology, progression, differentiation,. (2021).

9 de Bézieux, H. R., Van den Berge, K., Street, K. & Dudoit, S. Trajectory inference across multiple conditions with condiments: differential topology, progression, differentiation, and expression. *bioRxiv*, 2021.2003.2009.433671, doi:10.1101/2021.03.09.433671 (2021).

### Tables

**Table S1**: List of antibodies used for immunoblotting

| Antibody | Company* | cat number | dilution | Application** |
|---|---|---|---|---|
| Akt1 | CST | 2938S | 1:2000 | WB |
| Akt2 | CST | 2964S | 1:2000 | WB |
| AUKB1 | CST | 3094T | 1:2000 | WB |
| Delta N p63 (∆Np63-α) | CST | 67825S | 1:2000 | WB |
| ERK1/2 | CST | 4695T | 1:2000 | WB |
| FOXM1 | CST | 5436T | 1:2000 | WB |
| GAPDH-HRP conjugated | CST | 3683S | 1:2000 | WB |
| HMGB2 | CST | 14163T | 1:2000 | WB |
| Integrin α6 | CST | 3750S | 1:2000 | WB |
| Integrin β1 | CST | 34971S | 1:2000 | WB |
| K10 | Biolegend | PRB-159P | 1:2000 | WB |
| K14 | Serotech | MCA890 | 1:2000 | WB |
| K18 | CST | 4548T | 1:2000 | WB |
| K19 | Serotech | 5552-9009 | 1:2000 | WB |
| K7 | CST | 4465T | 1:2000 | WB |
| KLF4 | CST | 12173S | 1:2000 | WB |
| KLF5 | CST | 51586S | 1:2000 | WB |
| Myosin light chain 2 | CST | 8505S | 1:2000 | WB |
| p-Akt1 (S473) | CST | 9018S | 1:2000 | WB |
| p-Akt2 (S474) | CST | 8599S | 1:2000 | WB |
| p-c-Raf (Ser338) | CST | 9427T | 1:2000 | WB |
| pERK1/2 (Thr202/Tyr204) | CST | 4370T | 1:2000 | WB |

| | | | | |
|---|---|---|---|---|
| pMEK1/2 (Ser217/221) | CST | 9154T | 1:2000 | WB |
| p-Myosin light chain 2 (S19) | CST | 3675S | 1:2000 | WB |
| pRSK (Ser380) | CST | 11989T | 1:2000 | WB |
| Delta N p63 (∆Np63-α) | CST | 67825S | 1:50 | IF |
| K10 | Biolegend | PRB-159P | 1:500 | IF |
| Involucrin | Sigma | I9018 | 1:5,000 | IF |
| Filaggrin (AKH1) | Santa Cruz | 66192 | 1:500 | IF |
| K14 | Serotech | MCA890 | 1:500 | ICF |

*CST = Cell signalling Technology

**WB = western blotting; IF = Immunofluorescence;  ICF = Immunocytofluorescence

### Figures

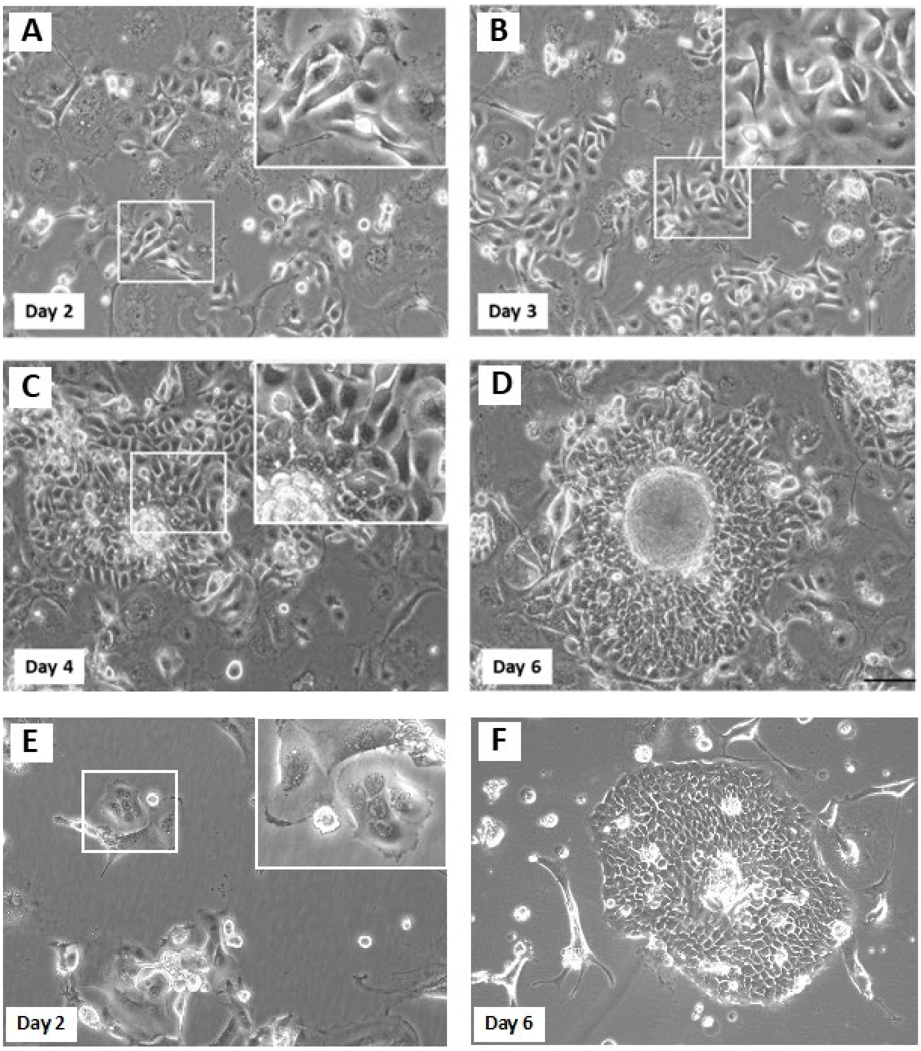


**Figure S1. Cell morphology and colony formation.**

Cells were treated with ROCKi and images were taken on days 2, 3, 4, and 6 (**A**–**D**). Images of the control cells on days 2 and 6 (**E** and **F**). Insets are the magnified regions indicated in white box. Scale bar = 100μM. Cells treated with ROCKi were dispersed on days 2 and 3, whereas control cells were already forming colonies on day 2.

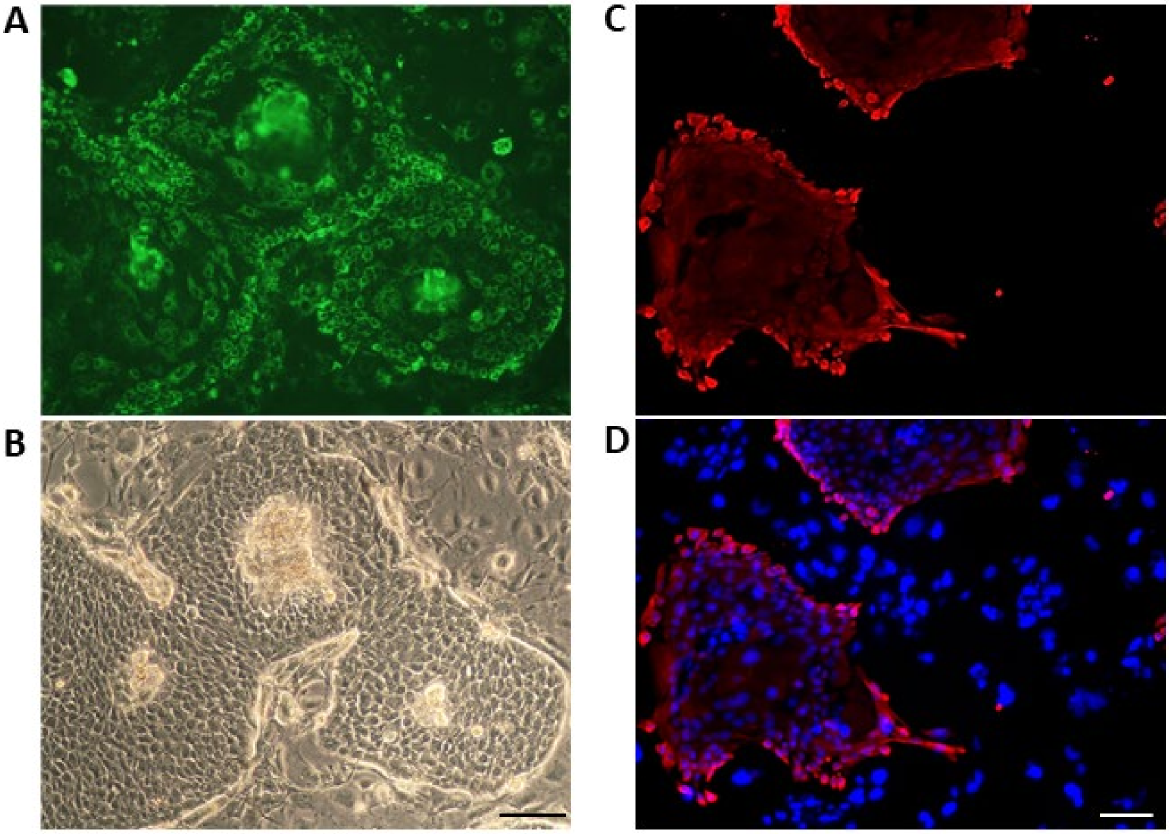


**Figure S2. Cyto-staining for mitochondrial mass and K14.**

Keratinocyte colonies six days after seeding were stained for mitochondrial content and K14 using MitoTracker Green and K14 antibody (**Table S1**). Mitochondrial content staining (green) and corresponding phase contrast images (**A** and **B**). Fluorescent immunostaining of K14 (red) (**C** and **D**). Nuclei were counterstained with DAPI. The distribution of both mitochondrial content and K14 were at the periphery of the colonies. Scale bar = 100μM.

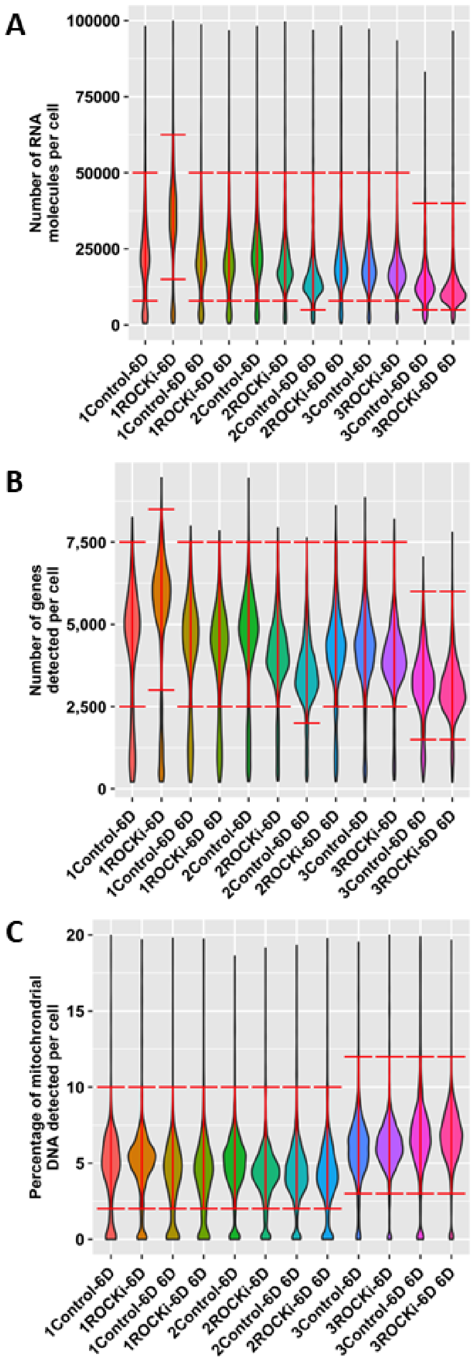


**Figure S3. Quality control (QC) thresholds.**

Violin plots for each quality control metric with the selected thresholds indicated by the red bars. A cell must lie within the thresholds for all three metrics to be used in the analysis. The number prepending the sample name corresponds to the donor of origin.

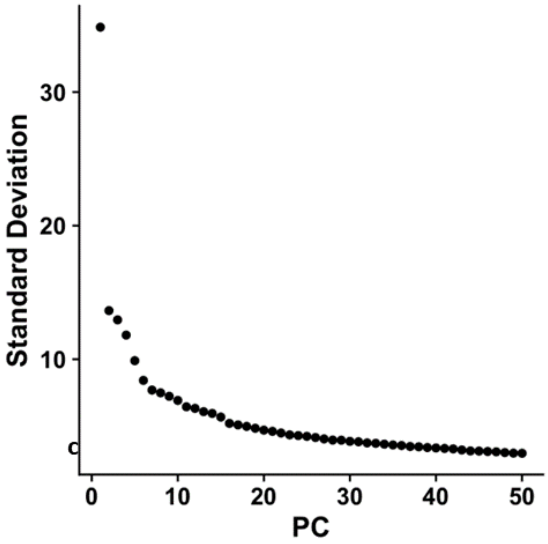


**Figure S4. An elbow plot showing the standard deviation of the first 50 principal components**.

There is no clear cut-off at which components appear to become less important except after the first five, which would have been too few. For this reason, we chose to use all 50 components.
